# Data-driven feedback augments ultrasound nanotheranostics in brain tumors

**DOI:** 10.1101/2025.05.01.651328

**Authors:** Hohyun Lee, Victor Menezes, Shiqin Zeng, Chulyong Kim, Cynthia M. Baseman, Jae Hyun Kim, Samhita Padmanabhan, Pranav Premdas, Naima Djeddar, Anton Bryksin, Nikhil Pandey, Pavlos Anastasiadis, Anthony J. Kim, Tobey J. MacDonald, Chetan Bettegowda, Graeme F. Woodworth, Felix J. Herrmann, Costas Arvanitis

## Abstract

The blood-brain barrier (BBB) renders the delivery of nanomedicine in the brain ineffective and the detection of circulating disease-related DNA from the brain unreliable. Here, we show that the acoustic emission content of focused ultrasound-controlled microbubble dynamics (MB-FUS) incorporates precursor signals that allow large-data models to predict sonication regimens for safe and effective BBB opening. Crucially, closed-loop MB-FUS controller augmented by machine learning (ML-CL) expands the treatment window (4-fold), as compared to conventional controllers, by persistently and proactively maximizing the BBB permeability while preventing tissue damage. By successfully scaling up from mice to rats and from healthy to diseased brains (glioma), ML-CL rendered the BBB permeable to large nanoparticles and markedly improved the release and detection of tumor DNA in plasma. Together, our findings reveal the potential of data-driven feedback to support the development of next-generation AI-powered ultrasound systems for safe, robust, and efficient nanotheranostic targeting of brain diseases.

## Introduction

The blood-brain barrier (BBB) constitutes a highly selective semi-permeable layer between the vascular lumen of capillaries and the brain parenchyma that tightly regulates the transport of macromolecules to support the metabolic demands of the brain and the clearance of soluble waste products ^1–4^. Due to its properties, the BBB hinders the effective delivery of a wide range of therapeutic agents (larger than 400 Da^5,6^) into the brain, including highly potent nanotherapeutics ^6–9^. It also limits the reliable release of soluble biomolecules into circulation – molecules that are associated with disease formation and progression – and are therefore essential for minimally invasive diagnosis and longitudinal monitoring of brain diseases through liquid biopsy techniques ^10,11^. Hence, strategies to safely and effectively overcome the BBB – including its partially compromised, yet highly heterogenous permeability during disease progression ^6,12^ – are critical for the diagnosis, treatment, and monitoring of central nervous system (CNS) diseases.

Circulating microbubbles (MB) upon focused ultrasound (FUS) exposure (sonication) can transiently increase the BBB permeability in a disease-agnostic way ^13–15^ and facilitate mass transport across this biological interface in an exposure-dependent manner ^16,17^. Notably, MB-FUS has been shown to increase the effective delivery of various therapeutic agents – including a broad spectrum of nano-formulations ^16^ – across a wide range of brain diseases ^16,18,19^. Despite promising preclinical findings, the quest to deliver an increasing fraction of intravenously administered nano-carriers into the brain ^20^ combined with the need to incorporate larger and more potent cargos (e.g., gene editing nano-systems) that inevitably require larger constructs (≥60 nm) ^21^, is probing scientists and clinicians to employ FUS exposures close to safety limits in order to maximize their penetration across the BBB ^22–27^. Similar advancements and challenges have also been observed in the use of MB-FUS to promote the release of disease-associated soluble molecules from the brain into circulation. The integration of MB-FUS with liquid biopsy techniques – whereby biomarkers from sonicated tissues are transiently enriched in the circulation – enables the development of novel assays for diagnosing and characterizing brain diseases ^28–35^. This includes the release of larger and scarcer molecules – such as circulating-tumor DNA (ctDNA, fragments ≥ 60 kDa) – which are critical for improving the sensitivity, specificity, and ultimately the diagnostic utility of these assays. However, such sono-interrogation approaches must carefully balance the need for aggressive sonication to improve biomarker release against the risk of inducing brain injury ^29–32,35^.

Balancing BBB opening efficacy with safety requires effective and real-time control of the cerebrovascular MB dynamics ^16^ – a highly nonlinear phenomenon that is prone to instabilities, such as MB collapse (i.e., inertial cavitation) that has long been linked to tissue damage ^36^. Recent investigations have shown that recording and analyzing the acoustic emissions (AE) generated by MBs’ nanoscale oscillations – which incorporate strong nonlinear acoustic components – provides a robust way to adjust the sonication settings ^16^. The sonication amplitude can be modified in real-time to attain prescribed strength of nonlinear acoustic components (i.e., Harmonics and/or Ultra-Harmonics) that are good proxies to stable oscillations and positively correlate with safe BBB opening. While this principle has formed the basis for developing real-time closed loop methods to maximize BBB opening ^17,37–46^, safety remains a concern as all current control methods are reactive to the onset of broadband emission: a warning sign associated with MB collapse. That is, they rely on the detection of broadband emission events before triggering a “safety response” (i.e., rapid pressure drop). As a result, small damage from repeated treatment sessions targeting neurologically functional brain regions (e.g., tumor invading healthy brain or neurological diseases like Alzheimer’s) can be compounded ^47–50^. Consequently, the “first do no harm” principle that dictates how new technologies and treatments are evaluated in the clinical space requires the adoption of conservative exposure settings during MB-FUS that inevitably leads to a very narrow acoustic treatment window (∼ 50 kPa) ^16,36^. The latter not only complicates the effective clinical translation and broad dissemination (i.e., similar to US imaging) of this technology, but also challenges its ability to consistently attain clinically meaningful endpoints related to the diagnosis, treatment, and treatment monitoring of CNS diseases.

Here, we employ data-driven models to extract precursor signals (i.e., patterns) ^51,52^ that are within the MB AE to study and analyze the transition from stable to inertial MB oscillation. Through comprehensive analysis and training of the models, using more than 54,000 AE datasets collected in MB-FUS experiments in mice, we discover multi-dimensional relationships in features derived from nonlinear MB AE that can predict the onset of broadband emissions with high sensitivity. By integrating the trained model into a real-time closed-loop MB AE feedback controller we show that it can augment the MB-FUS acoustic treatment window by persistently and proactively maximizing the BBB permeability while preventing tissue damage. We show that this data-driven feedback overcomes barriers to the delivery of nanoparticles and markedly improves the release of soluble biomarkers (protein and ctDNA) in orthotopic glioma tumor model in rats. Collectively, our findings demonstrate that, by expanding the treatment window, data-driven feedback can augment ultrasound nanotheranostic targeting of brain tumors and support the development of next generation AI-powered ultrasound systems for improved diagnosis, treatment, and monitoring of brain diseases.

## Results

### Multilayer perceptron (MLP) achieves high sensitivity in predicting broadband emissions

In this proof-of-concept investigation, we applied a supervised classification machine learning (ML) model trained on 54,040 acoustic emission (AE) datasets from BBB opening experiments across 415 distinct brain targets in 114 mice. All training data and experimental results presented in this study were acquired using a custom-built US-guided FUS system (USgFUS) operating at 0.5 MHz excitation frequency (see Methods) ^17^. As a proof-of-concept ML model, we employed a neural network model known as multilayer perceptron (MLP) consisting of fully connected neurons with one hidden layer (10 neurons) and a sigmoid activation function (**Fig. 1A**). Despite being a simple model, this structure allows the model to learn complex relationships between input dataset – MB AE, for instance – and broadband emission that are difficult to capture with conventional statistical tools ^53^. Moreover, when prior knowledge about feature relationships (i.e., MB AE vs. broadband emission) is limited or unavailable, MLPs – being a universal function approximators – offer a flexible solution compared to other ML models. We then trained the MLP model using Levenberg-Marquardt algorithm^54^. The inputs to the model included 12 features: primarily MB AE in the ultra-harmonic frequency range, which are considered to be closely related to inertial cavitation and brain damage ^40^, as well as additional variables such as MB kinetics and the presence of disease (cancer) in the targeted tissue (see **Table 1** for a complete list of features). Each dataset was labeled (i.e., ground truth labeling) based on whether a broadband emission event (defined as 6 dB above baseline noise) occurred in the subsequent sonication.

**Figure 1.**
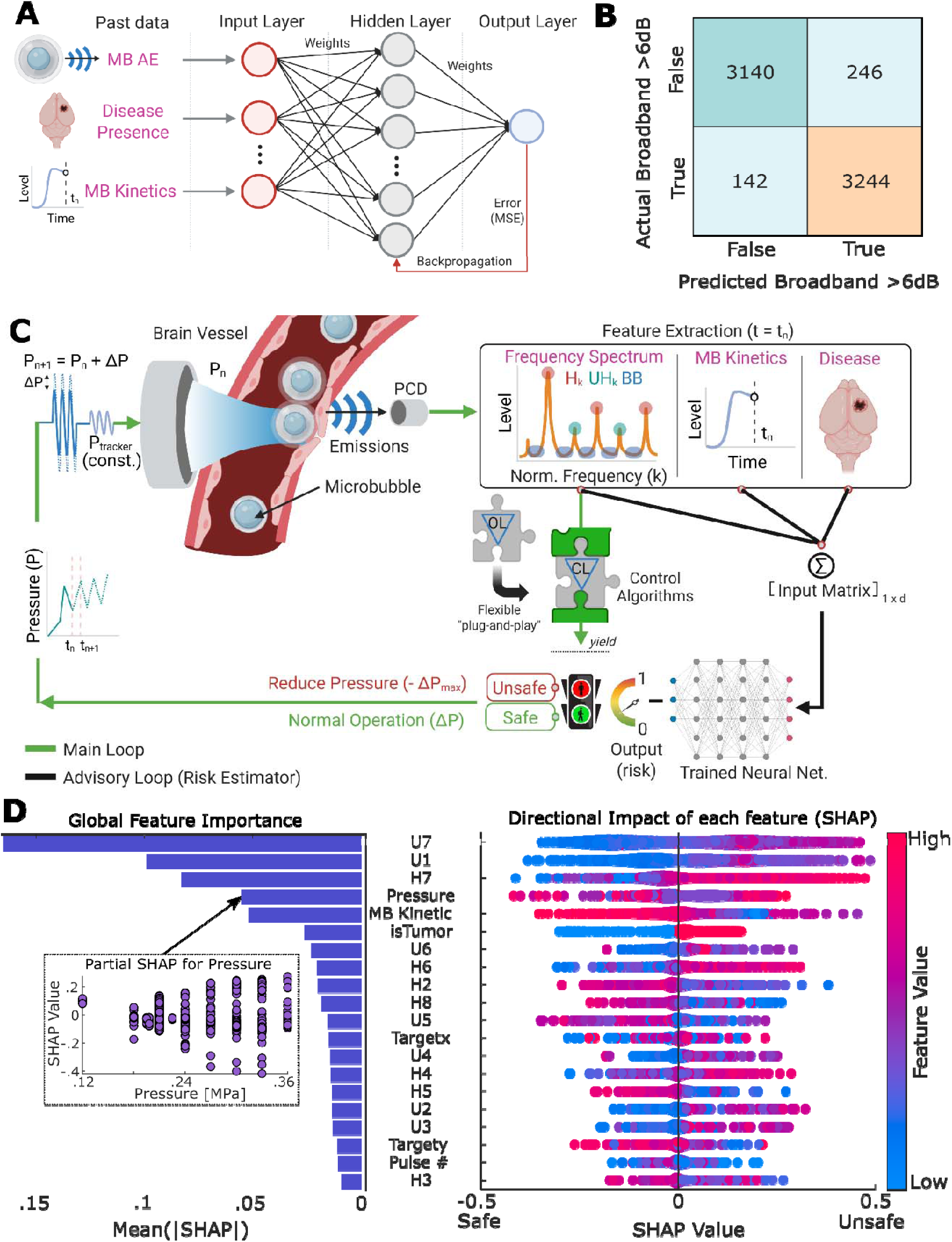
Multilayer perceptron model training, interpretation, and its integration with the controller. **A)** Schematic of simple proof-of-concept MLP model with one hidden layer with 10 neurons. Input features are described in **Table 1**. **B)** Confusion matrix for MLP model training (80% of ground truth label and matched number of ground false label); Numbers in the matrix indicate the number of data points with the top left, top right, bottom left, and bottom right being true negative, false positive, false negative, and true positive, respectively. **C)** Schematic of ML-assisted closed-loop MB-FUS controller. MLP model’s output is used as an additional safety layer (advisory loop) to override closed-loop controller’s decisions upon positive prediction. This flexible design allows its universal compatibility with various control algorithms. **D)** SHAP value analysis onto MLP model. All training datasets were used for analysis without random sampling. Left shows global importance calculated from the absolute value of SHAP values. Partial SHAP analysis of single feature, pressure, shows weakly positive correlation between pressure and SHAP, but with high variance, which indicates the effect of pressure onto model is not monotonic. For the right (SHAP analysis), being closer to red indicates higher values of each feature. SHAP value (x-axis) represents the contribution of each feature’s value onto the output of the model (positive SHAP value indicates the tendency of positive broadband prediction by the model).

**Table 1-.**
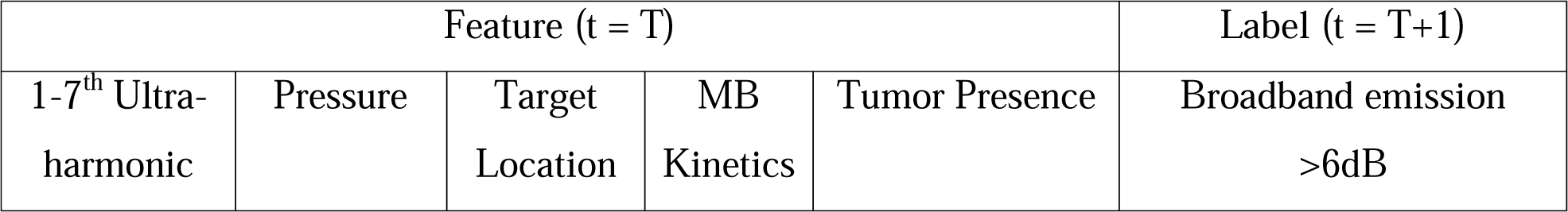
Features and Label for MLP model training.

Due to the inherent class imbalance (only 8% of total AE data contained broadband emissions) and to prevent model from being biased towards the more frequent no-broadband events, we performed under-sampling ^55^ of the majority class (i.e., no-broadband), which in fact showed more conservative behavior (i.e., higher false positive) while attaining sensitivity (i.e., true positive rate – correct broadband emission prediction) compared to oversampling technique (**Fig. S1 & Fig. S2**). We then trained the MLP model using 80% of the under-sampled data and tested it on the remaining 20% (see Methods). In our training dataset, we found 94% (6,384/6,772) overall accuracy, 96% (3,244/3,386) sensitivity (true positive rate), and 93% (3,244/3,490) precision (false positive vs. true positive), and 93% (3,140/3,386) specificity (true negative rate) in predicting broadband emission events (**Fig. 1B**). Crucially, the retained high sensitivity in testing dataset suggests that the model remains effective at detecting true broadband emission events, which is especially important given their scarcity and potential safety implications. The low precision in testing data set (53%, **Fig. S1**) reflects a tendency toward false positive predictions – possibly influenced by the simplicity of our proof-of-concept MLP model. On the other hand, low precision highlights the model’s conservative behavior, which is a desirable trait in the context of clinical translation, where minimizing the risk of missing predictions is a priority. We also confirmed that other machine learning models, such as support vector machine (SVM), had comparable performance (**Fig. S1 & Fig. S2**). Importantly, we found that when it is necessary to improve the performance and precision (i.e., reduce conservativeness) of the model, more advanced deep learning techniques such as the Attentive Multilayer Perceptron (AMP) model are highly potent (**Fig. S3 & Fig. S4**, see methods and Supplementary Information for detailed design). Together, these findings highlight the potential of ML methods to identify precursor patterns of future MB collapse. By anticipating rather than reacting to these adverse events, such data-driven methods offer the potential for real-time safety monitoring during MB-FUS treatment (i.e., responding before MB collapse and associated brain damage has taken place).

### Design and interpretation of machine learning-assisted closed-loop controller (ML-CL)

#### Design of machine learning-assisted closed-loop controller framework

Having established the predictive potential of the MLP model, we then designed a machine learning-assisted closed-loop MB-FUS controller (ML-CL) by integrating the trained MLP model into a closed-loop controller (CL) algorithm we have recently proposed and validated ^17^. CL algorithms operate by targeting a desired AE level (i.e., target level) and dynamically adjusting their input pressure based on real-time feedback from the current AE level, with the goal of minimizing the error between the current and target AE levels ^16^. In this framework, on top of the CL algorithm, we implemented the MLP as an additional safety layer (**Fig. 1C**). This design preserves the controller’s responsiveness to broadband emission events (i.e., reactive element) while introducing an additional predictive element that enables the controller to also anticipate broadband emissions and respond as if they have already occurred. Importantly, integrating MLP as an additional safety layer does not interfere with the controller’s actions (i.e., controlling harmonic emissions) during normal operation but ensures that the MLP acts when necessary – overriding the controller’s decision upon prediction of a broadband event during sonication. Perhaps the most salient feature of this design is its flexibility, as it allows the MLP to be integrated into any controller or system (provided there is sufficient AE data for training from ongoing trials or experiments) without altering its core functionality or features. Likewise, different ML models (see Supplementary Information) can be integrated into this framework.

#### Interpreting machine learning models offer a data-driven approach for studying and analyzing the MB dynamics in brain vessels

In light of the above findings, we took this analysis one step further by attempting to interpret the trained MLP model and support a data-driven analysis of MB dynamics. To interpret our MLP model, we assessed our results using Shapley Additive Explanations (SHAP) (**Fig. 1D**). SHAP assigns each input feature a contribution score (SHAP Values), indicating how much it influences the model’s output ^56^ and hence enabling quantification of their importance. While the original MLP model was trained using ultra-harmonics as the MB AE features, we included both ultra-harmonics and harmonics in the SHAP analysis to evaluate the broader importance of MB AE components (see **Table 2** for a complete list of features). As indicated by SHAP values, we found that the top 7 contributing input features were strongly associated with the key physical characteristics of the MB dynamics and the configuration of the USgFUS system (**Fig. 1D**). These were: i) the passive cavitation detector’s (PCD) peak sensitivity at 3.5 MHz, represented by 6^th^, 7^th^ ultra-harmonic and 7^th^ harmonic; ii) the MB dynamics strength and proximity to MB instabilities (i.e., bubble collapse), represented by 1^st^ ultra-harmonic, which are unique signatures of MB dynamics and have been used in the past to control the MB dynamics and establish safety thresholds ^40,42^; iii) the number of MBs in focal region, represented by MB kinetics; and iv) the peak negative pressure, which is a key driver of MB dynamics ^57^. Interestingly, while lower-order harmonics are often considered more useful in practice – owing to their efficient emissions ^58^ and effective transmission through the skull – our results suggest that the harmonics close to the receiver sensitivity (even up to 7^th^ harmonic) exhibit similar significance. This highlights the importance of the system’s design and tuning for recording MB AE. In our preclinical system, harmonics near 3.5 MHz (7^th^ harmonic) were shown as critical, which suggests that controlling 7^th^ harmonic (3.5 MHz) should be prioritized with this system configuration. We anticipate that this system-specific result will be different for clinical systems, however our framework can be applied as is using data from such systems. Our analysis also revealed that the presence of tumors can elicit higher probability of broadband emissions, which is surprising as this MB behavior has not been reported in the past despite extended preclinical and clinical investigations.

**Table 2-.** Features and Label for MLP SHAP Analysis.

| Feature (t = T) |  |  |  |  |  | Label (t = T+1) |
| --- | --- | --- | --- | --- | --- | --- |
| 2-8<sup>th</sup> Harmonic | 1-7<sup>th</sup> Ultra-harmonic | Pressure | Target Location | MB Kinetics | Tumor Presence | Broadband emission >6dB |

While most features showed consistent behavior – with high (red) or low (blue) feature values pushing the model’s predictions in a clear direction, **Fig.1D** – SHAP analysis revealed a more complex pattern for sonication pressure. We found that while pressure was a highly influential variable, its effect on the model’s output was not monotonic. This suggests that although sonication pressure plays a key role in shaping MB dynamics, its relationship with broadband emission is likely situation dependent. That is, despite a weak positive overall correlation (**Fig 1D**) between sonication pressure and broadband emission, this effect can be influenced by the skull and (transient) MBs’ concentration-dependent dynamics, among others ^59,60^. Moreover, by analyzing our training dataset, we found a close correlation between MB kinetics (5^th^ feature of importance according to MLP’s SHAP values) and the strength of broadband emission, which suggested that stronger broadband emission likely occurs at higher (>70%) MB (transient) concentrations in the body after a bolus administration (**Fig. S5**). This indicates that the temporal MB dose fluctuation, in addition to the total MB dose ^60–63^, is an important safety consideration. Together with SHAP analysis, these findings underscore the inherent risk associated with the use of constant pressure sonications (currently widely used in clinics) and further support the use of “ramping up” (closed-loop) control algorithms ^17^. In aggregate, our results demonstrated the capability of ML model and data-driven analysis of MB dynamics to predict instabilities in their behavior (i.e., MB collapse) and uncover subtle but potentially important features that shape their highly nonlinear response *in vivo*.

### Safety and efficacy evaluation of machine learning-assisted closed-loop FUS controller (ML-CL) in BBB opening

#### Evaluation of ML-CL at low exposure condition (32 dB)

To evaluate the operation of the ML-assisted closed-loop controller (ML-CL), we first performed MB-FUS mediated BBB opening in healthy mice by targeting three regions in each brain hemisphere using the USgFUS system that we used to collect the training data. We selected a target level of 32 dB at 7^th^ harmonic emission using the cavitation threshold curve identified from our training dataset (**Fig. S6A**). We chose this target level because it was at the highest level within the linear (before plateauing) regime of the curve. With 32 dB target level, we found that the average and maximum pressure decisions (P_Avg_ and P_Max_, reported as peak negative pressures) of ML-CL were 0.17 MPa and 0.23 MPa, respectively (**Fig. S6B**). Along with the pressure corresponding to 32 dB in the cavitation threshold model (P_Model_ = 0.14 MPa, **Fig. S6A**), these pressures provided the necessary exposure conditions for implementing constant pressure open-loop controllers (OL) to compare and benchmark the ML-CL controller’s performance. Unfortunately, the selected target levels and exposure settings were characterized by a low incidence of broadband emission events (3 events per 1,800 pulses) and, as such, they did not allow us to fully expose their potential benefits of ML-CL and tradeoffs between safety and efficacy (see below and **Fig S6**). Despite their shortcomings, they allowed us to establish feasibility for the proposed methodology in predicting and preventing scarce broadband emission events (See suppl. methods and **Fig S6**). They also provided an upper safety limit for the conventional (i.e. reactive) control methods as well as a lower BBB opening limit (P^32dB^_Avg_).

#### Evaluation of ML-CL at high exposure condition (36 dB)

In subsequent experiments (**Fig. 2A**), we selected a 7^th^ harmonic target level of 36 dB, which lies beyond the range of the cavitation threshold curve. The absence of representative pressure from the cavitation threshold model (P_Model_ = N/A) indicates that the 36 dB target level presents a challenging condition for all controllers (**Fig. 2B**). Note that while the threshold selected is above the controller calibration curve, it is still within the distribution of the data that we used to train the MLP, which is critical for accurate predictions (i.e., no extrapolating out of distribution). Again, the OL controllers operated at the pressure levels determined by the ML-CL operation (at 36 dB target level). That was at average (P_Avg_) and maximum pressures (P_Max_) of 0.25 MPa and 0.33 MPa, respectively (**Fig. 2C)**. We also incorporated a state of the art closed-loop controller (CL)^17^, operated at 36 dB target level. For each controller algorithm, we assessed the resulting AE, total broadband emission events, and BBB opening strength, as before (**Fig. 2D-E**). While in this condition, we found no significant difference in AE levels achieved among controllers (i.e., presumably due to AE saturation as observed in **Fig. 2B**), closed-loop control algorithms CL and ML-CL exhibited significantly lower AE fluctuation compared to the OL throughout the sonication (**Fig. 2F**), which reflects their ability to consistently maintain certain level of MB dynamics. Similar to AE, K_trans_ values indicated comparable BBB permeability across all groups (**Fig. 2G-H**), except for P_Avg_ and CL. The lack of significant difference between the groups is likely due to the high variance observed in the K_trans_ for OL group, which may stem from its corresponding variability in AE. Importantly, when we assessed the presence of broadband emission events, we found that OL controllers operated at P_Avg_ and P_Max_ resulted in 1.05% (11/1048) and 4.39% (46/1048) of sonications with broadband emissions, respectively (**Fig. 2I**, orange and red). Interestingly, CL group also showed a notable presence of broadband emission events in 1.34% (14/1048) of sonications, indicating that 36 dB target level not only represents a limiting condition for OL control algorithms but for all the controllers (including CL) that react to the broadband events (**Fig. 2I**, orange, red, and green). In contrast, the proposed ML-CL algorithm recorded only one instance of broadband emission event (1/1048, less than 0.1%), representing a 93% reduction compared to the CL algorithm (**Fig. 2I**, blue). This reduction is likely due to the MLP algorithm’s predictions during ML-CL operation, with 4.4% (46/1040) of its sonications flagged as potential broadband emission events (**Fig. 2J**). This is almost 20-fold higher prediction rate compared to 32 dB target level (**Fig. S6I**), highlighting the versatility of ML-CL and its ability to maximize the controller’s efficiency while maintaining a strong safety profile. Furthermore, applying MLP to the OL algorithms’ post-operation AE revealed that it could have predicted 51% of OL’s broadband emission events (**Fig. S7**).

**Figure 2.**
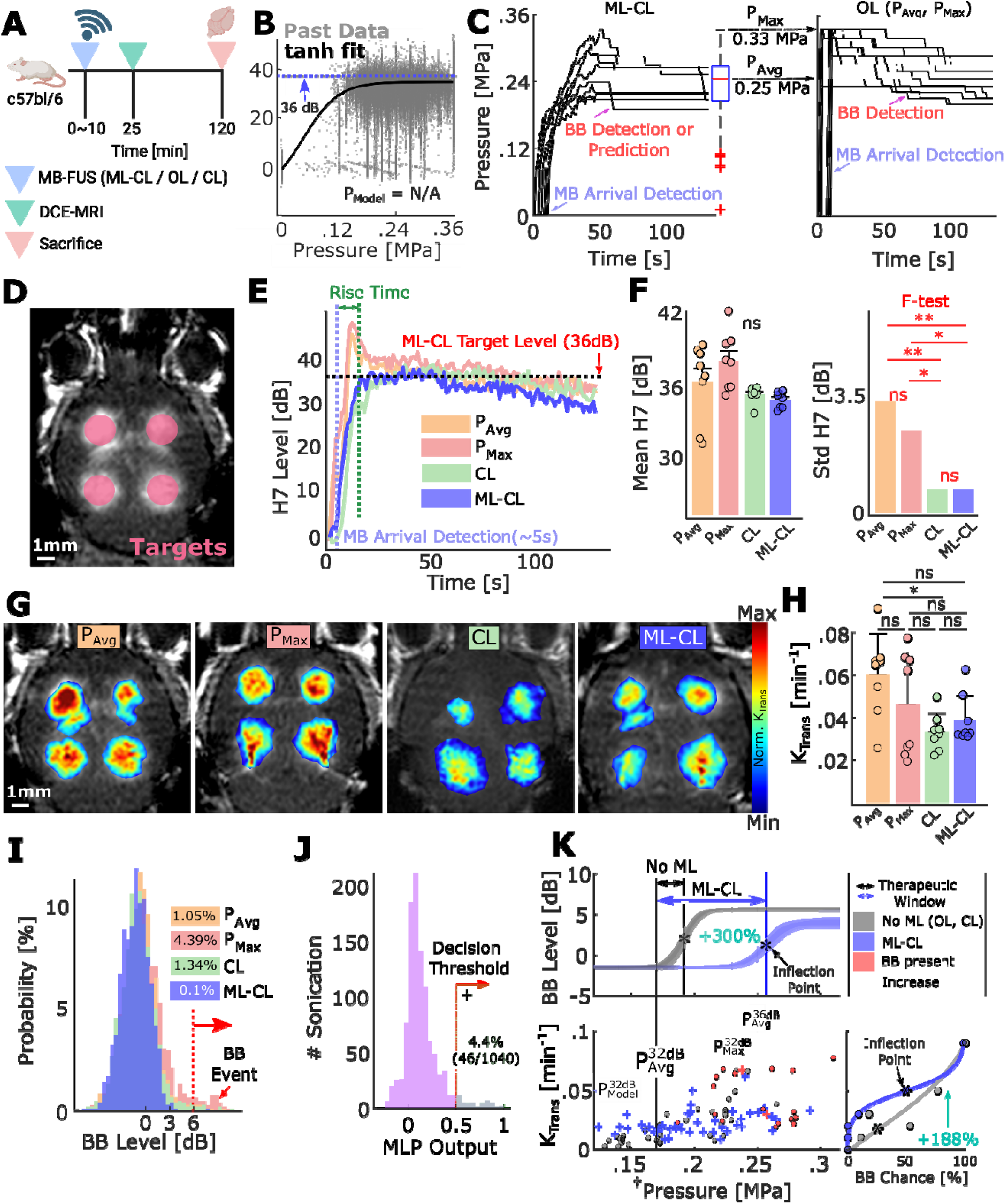
Performance assessment of ML-CL under limiting conditions. **A)** General MB-FUS experiment procedure for all controllers. **B)** Cavitation threshold model from the training dataset. 7^th^ harmonic AE as a function of pressure. To assess the controllers’ performance at their limits, we selected a target level above the model (36 dB). **C)** ML-CL pressure decisions for 36 dB target level (left). The average and maximum pressures used by ML-CL were used to determine P_Max_ and P_Avg_ for OL controllers (right). A decrease in pressure in OL indicates a reaction to broadband emission events (> 6 dB). **D)** Representative MRI image for 36 dB target level sonication targets. 2 targets in each brain hemisphere (total 4 targets per mouse) were treated. **E)** 7^th^ harmonic emissions during sonication for each controller. P_Avg_ : 36.2 ± 3.3 dB, P_Max_ 37.9 ± 2.4 dB, CL : 35.1 ± 0.7 dB, ML-CL : 34.7 ± 0.7 dB. n = 8 targets (total 2 animals) per group. **F)** Mean (left) and standard deviation (right) of 7^th^ harmonic emission during sonication by ML-CL. A variance test (f-test) was performed to compare standard deviation (right) of the mean 7^th^ harmonic emission at each target. **G)** MRI T1 images using the controller at 36 dB or equivalent OL pressure. **H)** Quantification of K_trans_ values through DCE-MRI. **I)** Histogram for broadband emission levels during sonication. Broadband emissions higher than 6 dB were considered a broadband emission event, whose probability for each controller is highlighted next to the dotted line. Percentages indicate 11, 46, 14, and 1 instances of broadband emission event out of 1,048 total sonications for P_Avg_, P_Max_, CL, and ML-CL, respectively. **J)** Histogram for MLP decision during ML-CL sonication. A model output of 0.5 or greater was considered a positive prediction. For this target level (36 dB), the model predicted 4.4% (46/1040) of sonication. **K)** Treatment window analysis for reactive (OL, CL) and predictive (ML-CL) controllers. Bottom left: mean pressure vs. K_trans_ at each target. Black indicates OL and CL, blue indicates ML-CL, and red indicates where broadband emissions were present. Top: pressure vs. broadband level throughout all sonication AE datasets. A more detailed figure with individual data points is shown in **Fig. S8**. ML-CL treatment window: 0.09 MPa (0.17 to 0.26 MPa). This was approximately a 300% increase (green area) from OL and CL’s treatment window: 0.02 MPa (0.17 to 0.19 MPa). Upper bounds for treatment windows were selected according to the inflection point of the hyper tangent curve fit, which indicates an increased likelihood of broadband emission. †The pressures (x-axis) used for the top left plot are the mean pressure at each targe brain location, and the pressure (x-axis) used for the bottom left plot is real-time pressure at each sonication similar to **B** (see **Fig. S8**). Also, for this figure, sonication data (K_trans_ and AE) from later parts of the study were included, which adds 1 additional broadband emission event for ML-CL. Bottom right: the probability of having broadband emission event vs. K_trans_. ML-CL improved K_trans_ given the same probability of broadband emission probability as evidenced by a shift in the curve by 188% (green area under hyper-tangent curve, the inflection point of 0.0173 to 0.0499). ns = not significant, *p<0.05, **p<0.01, ***p<0.001, and ****p<0.0001. Statistical analyses were performed through One-way ANOVA and Bonferroni correction. For the variance test, f-test was used with Bonferroni correction.

Subsequently, to determine treatment window improvement by ML-CL, we gathered the sonication data (e.g., pressure and broadband levels) from the reactive (OL, CL – i.e., No ML) and predictive (ML-CL) controllers and their resulting K_trans_ (**Fig. 2K**). We defined 0.17 MPa (P^32dB^_Avg_) as a conservative lower bound of the treatment window, as the K_trans_ rose above the background at this pressure (**Fig. S6G**). We then defined the upper bound of the treatment window as the pressure that resulted in a notable increase in broadband emission levels (i.e., point of inflection: 0.19 MPa for reactive controllers; 0.26 MPa for ML-CL). Strikingly, such pressure-bound analysis indicated that the ML-CL expanded the acoustic treatment window by 4-fold, from 0.02 MPa to 0.09 MPa (**Fig. 2K**, top). Moreover, by performing a similar analysis of K_trans_ and broadband emission event probability (**Fig. 2K**, bottom right), we found ML-CL achieved 3-fold increase in BBB permeability in healthy mice brains given the same risk of brain injury compared to reactive controllers (**Fig. 2K**, bottom right). Collectively, our results demonstrate that the integration of ML in MB-FUS mediated BBB opening control algorithms quadruples the treatment window, leading to improved BBB opening for a reduced risk of MB collapse (i.e., maximum benefit/risk ratio).

#### Safety confirmation of ML-CL at high exposure condition (36 dB)

Lastly, to directly assess the ability of ML-CL on preventing tissue damage, we examined the brain tissues sonicated with 36 dB target level using H&E staining (**Fig. 3A**). We found that all the MB-FUS target locations that were sonicated with the OL controllers (P_Avg_ or P_Max_) with broadband emission events had either visible or H&E-evidenced hemorrhages (i.e., petechiae) (**Fig. 3**). Similar responses were observed with the CL group although we identified petechiae at 2 (out of 8) targets where the broadband emission was undetected (**Fig. S9**). ML-CL group showed hemorrhage only at the one target where a broadband emission event was detected (**Fig. 3A**, bottom right). In a separate cohort, we sacrificed the animals 6 hours post sonication and stained the brains for astrocytosis (GFAP) and microglial activation (Iba-1), which are established markers for assessing neuroinflammation ^64–66^. We found that compared to the contralateral unsonicated region, P_Avg_ group showed an increase in GFAP expression (**Fig. 3C**, top), indicating early signs of astrocytosis in response to sonications. ML-CL revealed no significant increase in the GFAP expression, highlighting its ability to maintain safety. At this time point we did not observe differences in Iba-1 expression in either group. The observed lower expression of Iba-1 compared to GFAP expression is consistent with previous findings ^65^. Together our findings confirm that broadband emission events are key indicators of brain damage and demonstrate that integrating ML (i.e., predictive element) into a closed-loop controller can persistently and proactively ensure safety and consequently expand (4-fold) the treatment window of MB-FUS mediated BBB opening.

**Figure 3.**
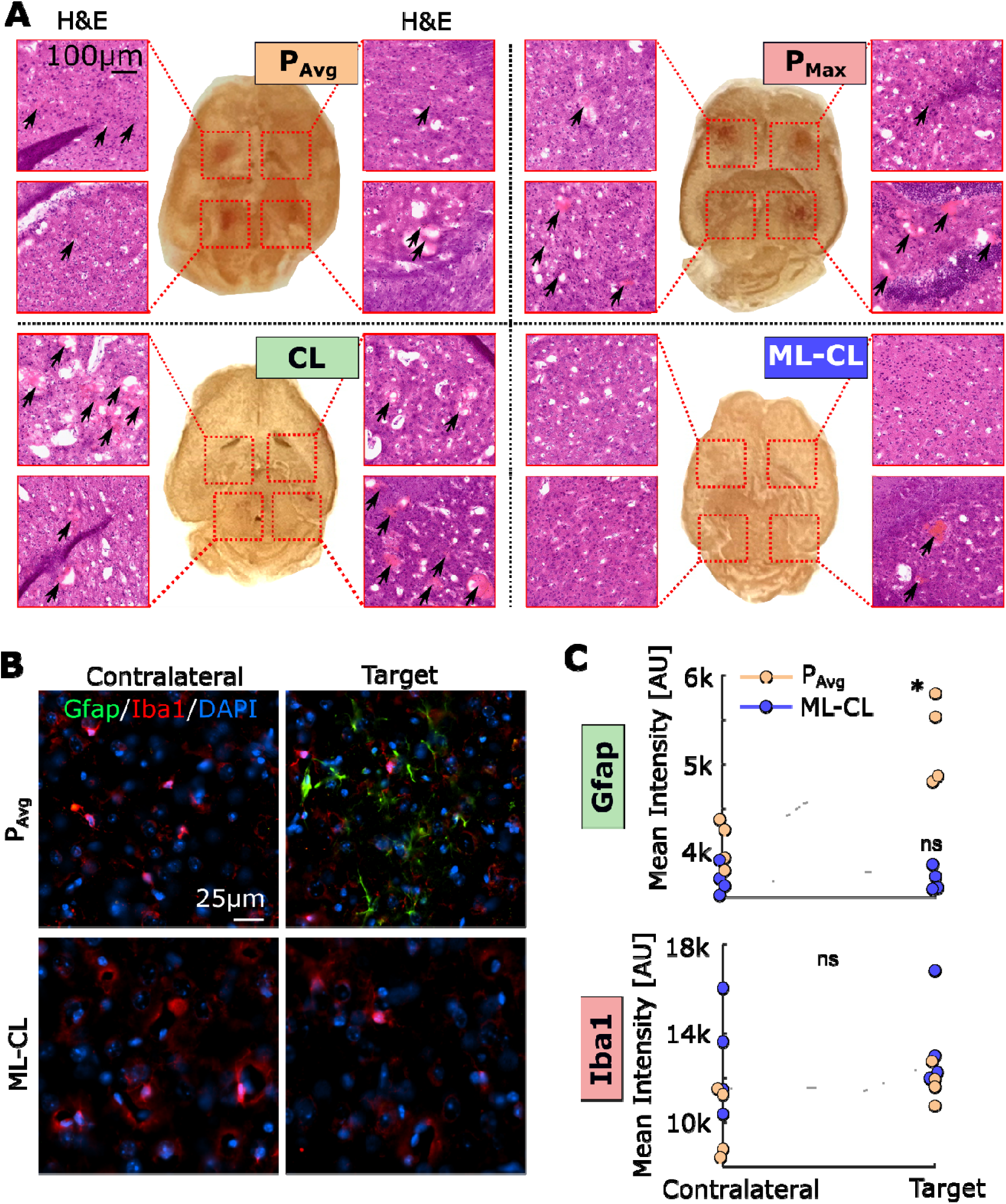
ML-CL can enhance safety without compromising BBB opening efficiency. **A)** H&E staining at each target referenced in photos of brain slices in the middle. **B)** Representative immunofluorescence images of GFAP and Iba-1. **C)** Quantification of GFAP and Iba-1 intensity comparing contralateral region and sonicated region. GFAP expression showed a significant increase (*p<0.05) in P_Avg_ group but not in ML-CL. No differences in Iba-1 were observed. Paired t-tests were used for statistical analysis. ns = not significant, *p<0.05.

### ML-CL augments the safe and efficient delivery of PEGylated polystyrene nanoparticles

Following the evaluation of ML-CL and establishing exposure conditions for robust BBB opening without compromising safety, we assessed the impact of the widened treatment window using ML-CL to deliver fluorescently labeled PEGylated polystyrene nanoparticles (NPs) of different sizes (37, 46, and 120 nm after PEGylation; detailed characterization shown in **Table 4**) ^67^. We sonicated healthy mice at one posterior and one inferior region on each brain hemisphere, where left and right hemispheres were sonicated with ML-CL operating at 32 dB and 36 dB, respectively (**Fig. 4A**). Immediately after sonication, we intravenously delivered NPs and analyzed the resulting delivery using IVIS (**Fig. 4B)**. During sonication (**Fig. S10 A-B**), we observed only one instance (1 out of 1048 total sonications) of broadband emission event at 36 dB target level (one target from the 120 nm group, **Fig. S10 C**). Hence, we deduced that the safety profile was comparable to our previous investigations (**Fig. 2 & Fig. 3**). Moreover, our analysis of K_trans_ (**Fig. 4C**) supported previous observations that indicated a target-level-dependent increase in BBB permeability in healthy brains. In agreement with the levels in AE and K_trans_, we observed enhanced NP delivery by 2.8-fold, 2.1-fold, and 2.1-fold for 37, 46, and 120 nm sizes, respectively (compared to non-sonicated regions – **Fig. 4D**; 36 dB targets). While the data had significant variation, presumably due to IVIS assessment (i.e., lower detection sensitivity for deeper targets), the NP delivery was significantly higher (p < 0.05) only for the 36 dB sonication. This indicates that the improved treatment window by ML-CL is critical for improving NP (up to 120 nm) delivery in the brain. We also investigated the size dependency of *in vivo* NP delivery by accounting for size-dependent total radiance (**Fig. 4E-F**). Interestingly, we found that 46 nm had maximum delivery (**Fig. 4G**). While these data seem counterintuitive, they support past studies that have shown that the delivery of NPs in the range of 40-50 nm is optimal after MB-FUS ^68^. In aggregate, these findings demonstrate the ability of ML-CL to significantly enhance the delivery of NPs in the murine brains without compromising safety by critically expanding the treatment window of MB-FUS.

**Figure 4.**
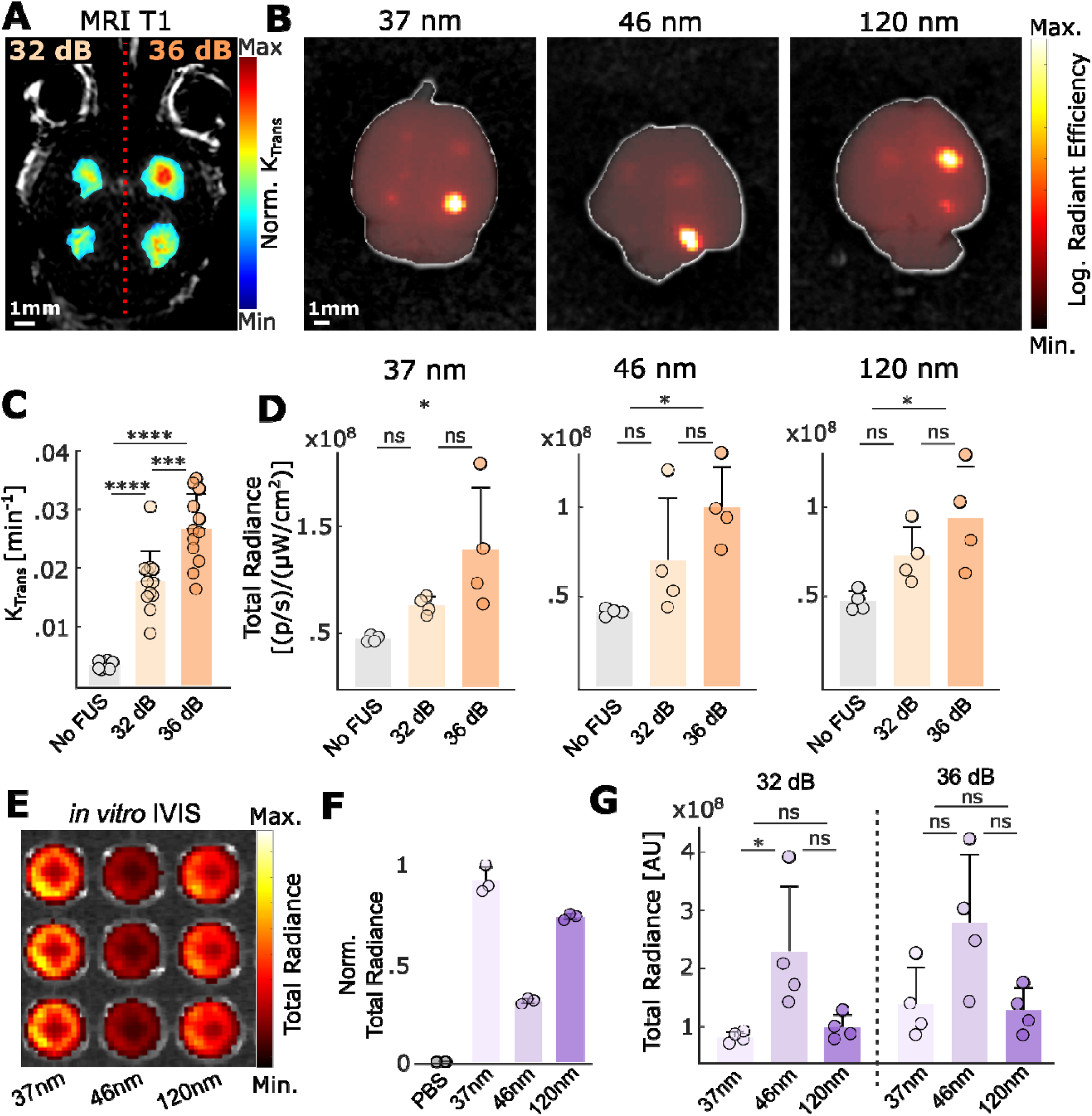
By expanding the treatment window ML-CL can safely enhance delivery of PEGylated polystyrene nanoparticles with broad size distribution in healthy brain. **A)** Representative T1-MRI image after ML-CL sonication. Left brain hemisphere was treated with 32 dB, and right brain was treated with 36 dB ML-CL. n = 2 animals (total 4 targets) per group. **B)** Representative IVIS imaging showing delivery of each size NPs at ML-CL targets. Color range is formulated in logarithmic scale. **C)** Quantification of K_trans_ for each target level. Both 32 and 36 dB target levels resulted in significantly higher K_trans_ compared to unsonicated region (p<0.0001). 36 dB group had higher K_trans_ compared to 32 dB group (p<0.001). Statistical analysis was performed through one-way ANOVA with Bonferroni correction. **D)** Quantification of IVIS fluorescence for each size NPs. 36 dB group increased 37, 46, and 120 nm NP delivery by 2.8, 2.1, and 2.1-fold, respectively (*p<0.05). Statistical analysis was performed using one-way ANOVA with Bonferroni’s correction. **E)** Representative image of *in vitro* IVIS setup with the different NPs (injection concentration) on a well plate (see Methods for details). **F)** We used the normalized *in vitro* radiance from each NPs to account for differences in their *in vivo* fluorescence. **G)** Normalized quantification of D using *in vitro* fluorescence ratio. ns = not significant, *p<0.05, **p<0.01, ***p<0.001, and ****p<0.0001. Statistical analyses were performed through One-way ANOVA and Bonferroni correction.

### ML-CL augments the release of brain cancer soluble biomarkers to the circulation

Subsequently, using tumor-bearing GL-261 mouse models, we assessed the ability of the ML-CL controller to augment the release of soluble biomarkers into the circulation. In this proof-of-concept investigation, we transduced the GL-261 cell line with Gaussia Luciferase (GLuc), so they can release a luminescent protein ^69,70^. This allows us to concurrently compare the release of GLuc protein (∼ 20kDa) and gene (∼ 60kDa) ^71,72^ with high specificity. We sonicated the GL-261-GLuc tumor-bearing mice using ML-CL (32 or 36 dB target level) and collected blood pre- and post-treatment (**Fig. 5A**). For non-sonicated control group (i.e., MB only; No FUS), we followed the same blood collection protocol. We chose 32 or 36 dB ML-CL target levels based on the NP delivery experiments, where the former represents a conservative exposure (i.e., close to current safety levels) and the latter can consistently improve the transport of large molecules across the BBB (**Fig. 4**). The tumor sizes were evenly distributed among the groups (**Fig. 5B-C**), and treatment groups were sonicated in 5 locations across the tumor; this sonication protocol (multiple-targets to cover tumor) was employed in all subsequent experiments involving tumor-bearing rodents.

**Figure 5.**
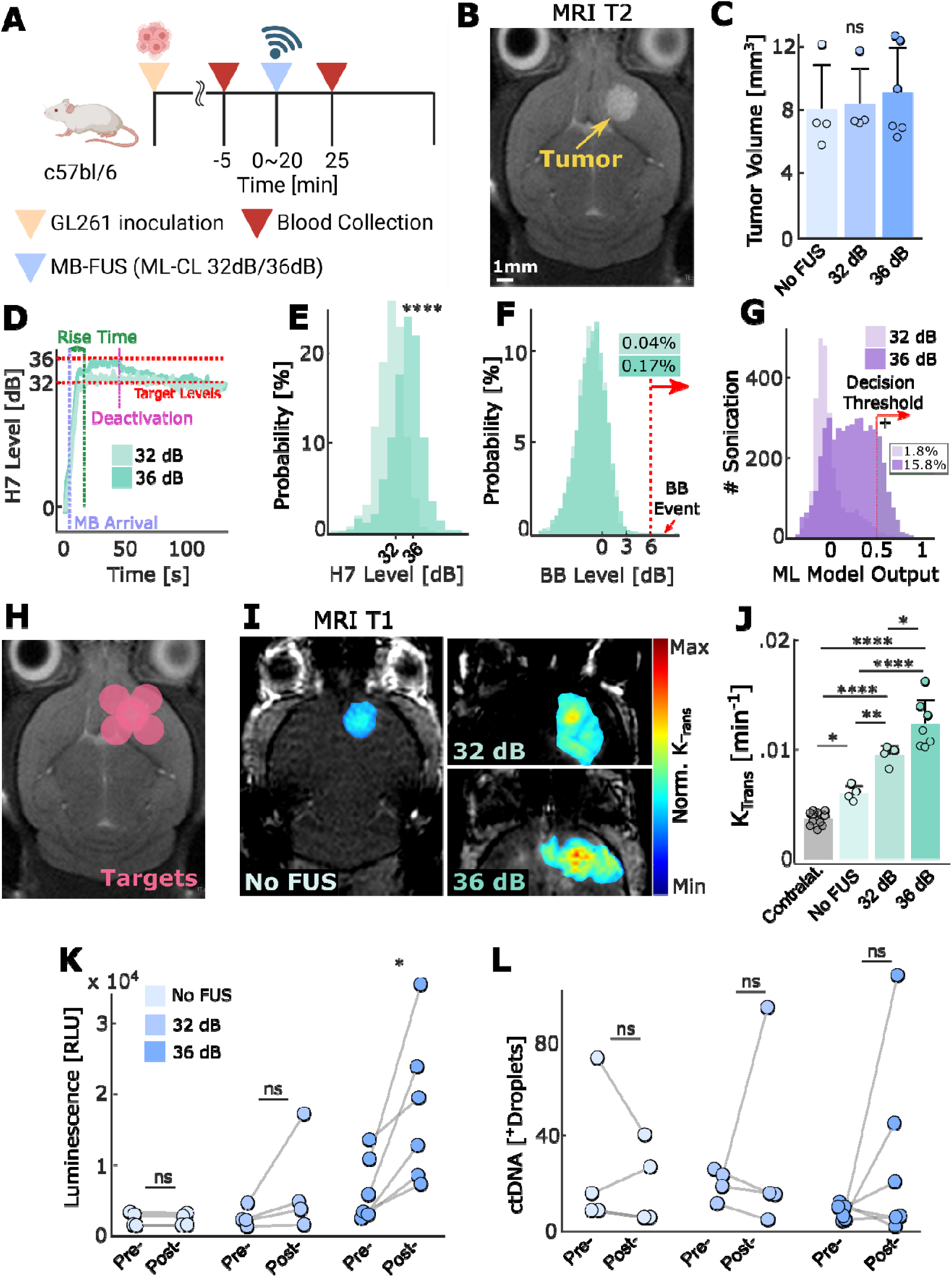
ML-CL augments the release of brain cancer-soluble biomarkers. **A)** Experimental procedure indicating the time points for treatment and blood collection. The control group was injected with MB only as treatment. **B)** Representative T2-MRI images for tumor, GL-261-GLuc. **C)** Tumor size distribution across groups. All groups had no significantly different tumor size distribution. (n = 4, control, n = 4, 32 dB, n = 6, 36 dB, respectively). **D)** Transient 7^th^ harmonic emission of 32dB (light green) and 36 dB (green) ML-CL during its application on GL-261-GLuc tumor-bearing mice. **E)** Distribution of 7^th^ harmonic emission for ML-CL at 32 dB and 36 dB target levels. 7^th^ harmonic level distribution was compared with t-test. (32 dB – 30.96 ± 3.13 dB; 36dB – 34.57 ± 3.73 dB). **F)** Histogram of broadband emission during sonication. 32 dB ML-CL had 0.04% (1/2620), and 36 dB had 0.17% (8/4585) of sonication that contained broadband events. G) ML model output during sonication. The model predicted broadband emission at 1.8% (48/2620) and 15.8% (722/4585) of 32dB and 36 dB ML-CL sonication, respectively. **H)** ML-CL target locations. **I)** Representative MRI-T1 images and **J)** quantification of K_trans_. **K)** GLuc protein luminescence quantification. Mean relative change – Control group: no difference (ns), 32dB: 2.2-fold (ns), 36 dB: 3.1-fold (p<0.05). **L)** GLuc gene quantification. Mean relative change – Control group: no difference (ns), 32dB: 1.5-fold (ns), 36 dB: 7.8-fold (ns). ns = not significant, *p<0.05, **p<0.01, ***p<0.001, ****p<0.0001. All statistical analyses were performed using One-way ANOVA with Bonferroni correction. Specific samples were excluded, check methods for exclusion criteria.

Interstingly, we found that the average harmonic target level achieved by ML-CL (34.57 ± 3.73 dB) was lower than in healthy brain study (35.1 ± 0.7 dB) (**Fig. 5D-E**). While this is somehow surprising, it is nevertheless consistent with our observations during SHAP analysis of the MLP, which suggested that the presence of a tumor drives the model towards positive predictions (**Fig. 1D**), which activates the MLP and forces the controller towards lower pressures and consequently towards lower harmonic levels. In fact, while we saw similar probability of broadband emissions compared to healthy brain (0.17% vs. 0.1%) (**Fig. 5F**), we also observed an increase in total predictions made by MLP (15.8%) in the presence of tumor as compared to healthy brain (4.4%) (**Fig. 5G**), further corroborating the SHAP analysis. These results highlight ML-CL’s versatility and robustness to different biological environments of tumor-bearing mice.

As evidenced by the increase in K_trans_, non-sonicated control tumors were leakier than contralateral healthy brains. However, after ML-CL operation on tumors (**Fig. 5H**), there was a significant increase in BBB permeability in sonicated tumors as compared to control group tumors (**Fig. 5I-J**). Crucially, we found that only for ML-CL operating at 36 dB, the GLuc protein concentration in the blood significantly increased (3.1-fold) immediately after sonication (p<0.05) (**Fig. 5K**). This suggests that, similar to therapeutic agent delivery, release of tumor-soluble macromolecules require the higher exposure conditions that only the ML-CL can attain without compromising safety. Meanwhile, although these GLuc protein levels persisted after 2 hours, a declining trend was evident (**Fig. S11**), suggesting a burst release immediately after BBB opening. In contrast to GLuc protein, GLuc gene analysis showed that the levels of positive droplets in the circulation were very low and inconsistent for all groups (**Fig. 5L**), indicating insufficient amount of DNA fragments in the collected blood. While the analysis of the smaller and more abundant GLuc protein consistently demonstrated the critical role of BBB permeability in restricting the release of cancer soluble biomarkers, the lack of GLuc gene in our analysis highlighted several challenges in addition to BBB permeability. These are possibly related to the degradation of ctDNA^71^ in the circulation, assay sensitivity, or sampling issues related to the relatively low blood volume collection limit allowed for mice. To address the impact of these factors on the detection limit and decipher the role of BBB permeability on ctDNA detection, we decided to conduct experiments in larger animal models (see below).

### ML-CL improves the delivery of nanoparticles in glioma tumors in rats

To assess the robustness and scalability of the ML-CL framework and its ability to widen the treatment window, we scaled it up to rats – an inherently more challenging model due to a thicker skull and different MB clearance kinetics – and compared its performance against an open-loop controller operated at P_Avg_. Following the experimental protocols similar to those used in mice (**Fig. 6A-C**), we sonicated F98 tumor-bearing (SRG) rats with 7 non-overlapping locations with either ML-CL or P_Avg_. For ML-CL, we conservatively chose a target level of 33.5 dB to account for thicker skulls, and a corresponding P_Avg_ of 0.29 MPa was determined from ML-CL operation (**Fig. 6B**). In these experiments, we were intravenously delivered 46 nm NPs at a dose of 5 mg/Kg after sonication. Note that the minimum operation time for controller (i.e., MB kinetics recording) was increased to 40 seconds as compared to 20 seconds in mice to accommodate for the faster MB kinetics in rats.

**Figure 6.**
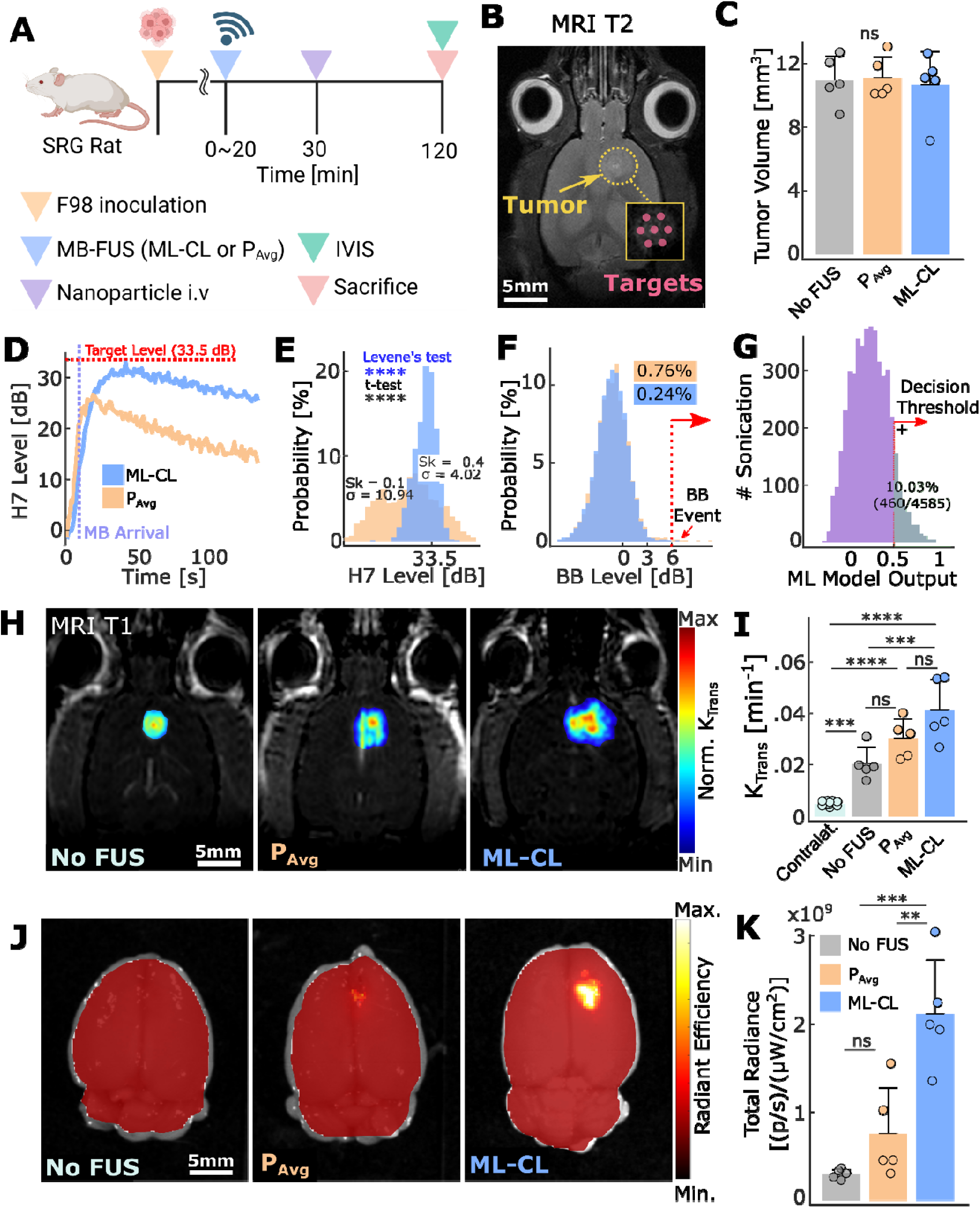
ML-CL augments delivery of PEGylated polystyrene nanoparticles in F98 tumor-bearing rats. **A)** Experimental procedure indicating the time points for treatment, NP administration, and IVIS analysis. The control group (no FUS) was injected with MB only as treatment. **B)** Representative MRI-T2 image for F98 tumor and sonication target locations. **C)** Tumor size distribution across groups. All groups had no significantly different tumor size distribution. (n = 5, control, n = 5, P_Avg_, n = 5, ML-CL 33.5 dB, respectively) **D)** Transient 7^th^ harmonic levels during treatment (Blue: ML-CL 33.5 dB; Orange: P_Avg_ 0.29MPa). Note that the curve is average of all targets, where MB arrival time and rise time can be different, that led to lower averaged harmonic level. **E)** Histogram of 7^th^ harmonic levels for P_Avg_ (Orange) and ML-CL 33.5 dB (Blue). 7^th^ harmonic levels during 20 seconds following MB arrival were considered for all sonication. 30.8 ± 4.0 dB for ML-CL; 24.7 ± 10.9 dB for P_Avg_. Levene’s test was used for variance comparison and t-test for mean comparison. Sk = skewness. **F)** Histogram of broadband emission during sonication. P_Avg_ (Orange) had 0.76% (35/4585), and ML-CL (Blue) had 0.24% (11/4585) of sonication that contained broadband events. **G)** ML model output during sonication. The model predicted broadband emission at 10.03% (460/4585) of ML-CL sonication. **H)** Representative MRI-T1 images and **I)** quantification of K_trans_. **J)** Representative IVIS image of NP delivery in F98 tumor-bearing rats. Left: Control (no FUS); Middle: P_Avg_ (0.29 MPa); Right: ML-CL (33.5 dB target level). **K)** Quantification of NP delivery. ML-CL improved NP delivery by 7.2-fold (p<0.001) compared to control group and 2.8-fold (p<0.01) compared to P_Avg_ group. ns = not significant, *p<0.05, **p<0.01, ***p<0.001, ****p<0.0001. All statistical analyses were performed with One-way ANOVA with Bonferroni correction.

Interestingly, we observed that compared to P_Avg_, ML-CL achieved a higher mean harmonic level (30.8 ± 4.0 dB vs. 24.7 ± 10.9 dB – **Fig. 6D-E**) and actively shaped its distribution, resulting in a mildly left-skewed profile (skewness = −0.4) with lower standard deviation (10.9 dB vs. 4.0 dB; p<0.0001, Levene’s test; **Fig. 6D-E**). This suggests that in contrast to P_Avg_, ML-CL overcomes weak therapeutic activity and modulates harmonic emissions clustering towards higher values with consistency, thereby highlighting its ability to effectively scale its performance and adapt to different (larger) animal models. Importantly, both controllers maintained comparable and low broadband emission probability (0.76 % vs. 0.24% for P_Avg_ and ML-CL, respectively; **Fig. 6F**). The MLP model exhibited consistent behavior in rats as it did in mice (**Fig. 6G**), demonstrating that ML-CL’s safety performance can be scaled to larger animal alongside its controlling capability.

Our analysis of BBB permeability using K_trans_ values corroborated our observations in harmonic level distribution: permeability increased progressively from the control group (untreated) to P_Avg_ group and was highest in the ML-CL group (**Fig. 6H-I**). Surprisingly, despite this trend in K_trans_ – which also revealed the leaky nature of untreated tumors compared to healthy brain region (**Fig. 6I**) – we observed a notable difference in actual NP delivery. Specifically, NP delivery was negligible in the control group but significantly enhanced only by ML-CL, achieving a 7.2-fold and 2.6-fold increased delivery compared to the control group and P_Avg_ (**Fig. 6J-K**), respectively. This not only suggests that AEs’ may better predict larger molecule delivery in brain tumors than K_trans_ (i.e., maps the small molecule gadolinium permeability) but also re-emphasizes the potential of ML-CL to reliably and safely maximize therapeutic outcomes. Collectively, these findings demonstrate that ML-CL can be scaled to larger animal models while maintaining its performance (i.e., maximizing the BBB permeability while preventing broadband events) and ability to improve NP delivery in brain tumors.

### ML-CL augments the release of ctDNA into circulation from glioma tumors in rats

Finally, using F98 tumor-bearing rat models, we evaluated the ability of ML-CL controller to improve the release of GLuc protein and gene into the circulation. Compared to mice, rats permit the collection of larger blood samples that allow us to assess its impact on the detection of GLuc protein and gene. In addition to increased blood volume, here we also employed a pre-amplification step via nested-PCR technique to further improve the sensitivity of the assay. Similar to our mouse study, we transduced the F98 cell line with Gaussia Luciferase (GLuc) and quantified its protein and gene levels in the blood. As before, we sonicated the F98-GLuc tumor-bearing rats using P_Avg_ and ML-CL and collected blood pre- and post-treatment (**Fig. 7A**). For non-sonicated control group, we injected MB only (No FUS) with same blood collection protocol. We maintained tumor-sonication protocol similar to NP delivery study (**Fig. 7B-C**). For ML-CL target level, we chose the same (33.5 dB) as in our NP delivery, where P_Avg_ was found to be 0.30 MPa. For these experiments we chose immunocompetent rat strain (CDF), as it represents a more challenging and clinically relevant model (i.e. rapid clearance and degradation of biomolecules in the circulation). The implementation of the immunocompetent rats leads to lower biomarker concentration baseline^73,74^ (pre-FUS), which demonstrates more realistic impact of FUS on molecule release (**Fig. S12**).

**Figure 7.**
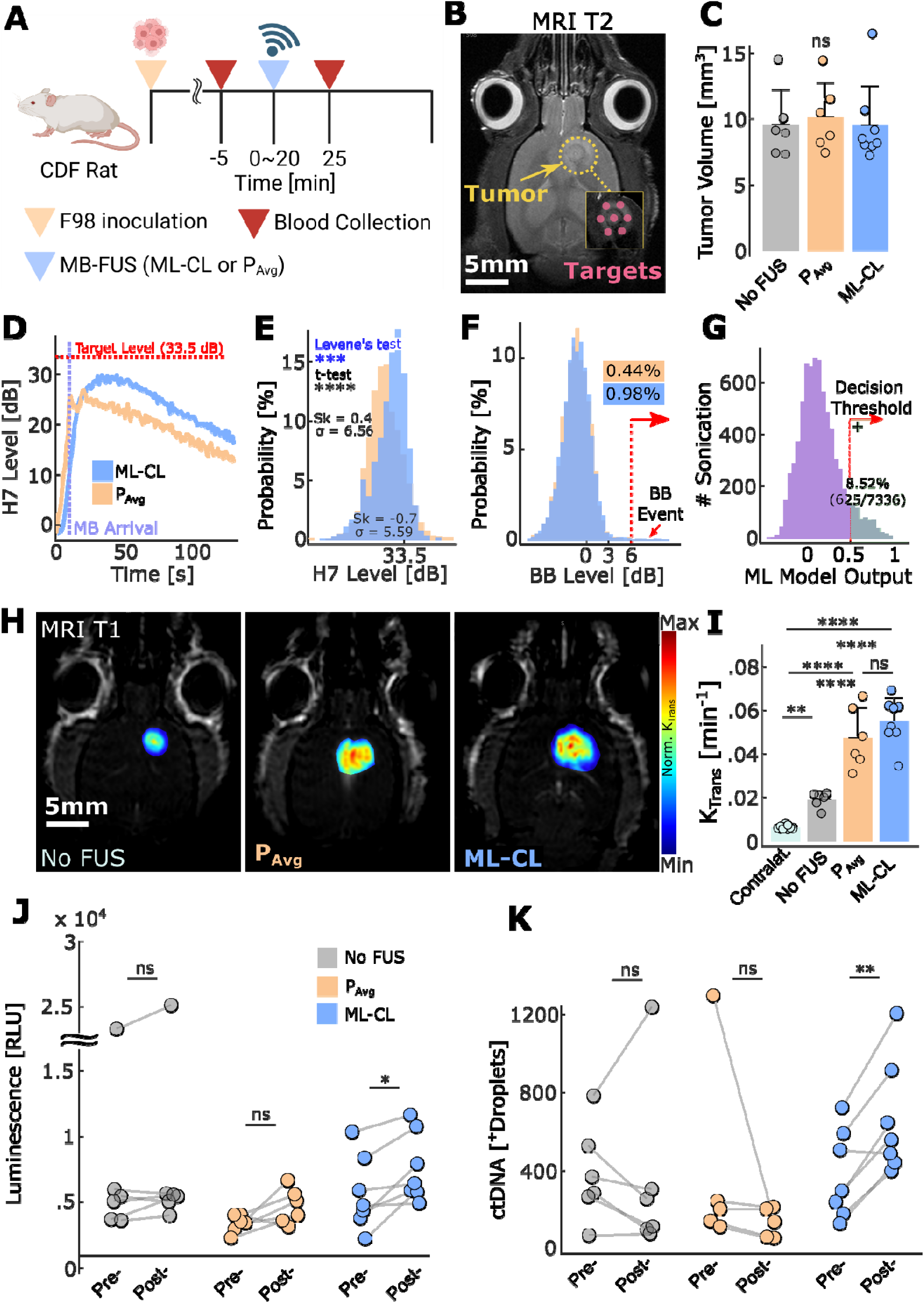
ML-CL augments release of protein and ctDNA from F98-GLuc tumor-bearing rats. **A)** Experimental procedure indicating the time points for treatment and blood collection. The control group (no FUS) was injected with MB only as treatment. **B)** Representative MRI-T2 image for F98-GLuc tumor and sonication target locations. **C)** Tumor size distribution across groups. All groups had no significantly different tumor size distribution. (n = 6, control, n = 6, P_Avg_, n = 8, ML-CL 33.5 dB, respectively) **D)** Transient 7^th^ harmonic levels during treatment (Blue: ML-CL 33.5 dB; Orange: P_Avg_ 0.30 MPa). Note that the curve is average of all targets, where MB arrival time and rise time can be different, that led to averaged down harmonic level. **E)** Histogram of 7^th^ harmonic levels for P_Avg_ (Orange) and ML-CL 33.5 dB (Blue). 7^th^ harmonic levels during 20 seconds following MB arrival were considered for all sonication. ML-CL: 28.2 ± 5.6 dB and P_Avg_: 26.8 ± 6.6 dB. Levene’s test was used for variance comparison and t-test for mean comparison. Sk = skewness. **F)** Histogram of broadband emission during sonication. P_Avg_ (Orange) had 0.44% (24/5502), and ML-CL (Blue) had 0.98% (72/7336) of sonication that contained broadband events. **G)** ML model output during sonication. The model predicted broadband emission at 8.52% (460/7336) of ML-CL sonication. **H)** Representative MRI-T1 images and **I)** quantification of K_trans_. **J)** GLuc protein luminescence quantification. Mean relative change – Control group: 1.1-fold (ns), P_Avg_: 1.4-fold (ns), 36 dB: 1.5-fold (p<0.05). Left: Control (no FUS); Middle: P_Avg_ (0.30 MPa); Right: ML-CL (33.5 dB target level). **K)** GLuc ctDNA quantification. Mean relative change – Control group: no difference (ns), P_Avg_: no difference (ns), 36 dB: 2.1-fold (p<0.01). ns = not significant, *p<0.05, **p<0.01, ***p<0.001, ****p<0.0001. All statistical analyses were performed with One-way ANOVA with Bonferroni correction. Specific samples were excluded, check methods for exclusion criteria.

Consistent with our previous findings, compared to P_Avg_, ML-CL produced a left-skewed (skewness = −0.7 vs. 0.41; **Fig. 7D-E**) and tighter (5.6 vs. 6.6 dB standard deviation; p<0.001; Levene’s test) harmonic distribution with higher mean (28.2 ± 5.6 vs. 26.8 ± 6.6 dB dB), reflecting its ability to control and maintain elevated and consistent harmonic levels, as compared to P_Avg_. Interestingly, unlike in the NP delivery study, P_Avg_ showed reduced harmonic emission variance, presumably due to smaller size and decreased skull thickness of this rat strain. Nonetheless, our results highlight the inherent shortcomings of open-loop (constant-pressure) controllers which are prone to instabilities in their harmonic emission such as i) right-skewed distribution – likely due to the early influx of MBs after bolus injection – that by nature contributes to lower mean and ii) higher variability in distribution. Importantly, given that both P_Avg_ and ML-CL showed less than 1% chance of broadband emission event, our results demonstrated that ML-CL’s safety performance is also robust to biological variability introduced by different strains.

Our analysis of BBB permeability using K_trans_ values, however, did not reflect our observations in the harmonic level distribution: while permeability increased progressively from the control group (untreated) to P_Avg_ group, ML-CL group’s permeability was not significantly different from P_Avg_ (**Fig. 7H-I**). Surprisingly, despite this lack of trend in K_trans_ analysis, which quantifies the permeability of the much smaller MRI contrast agent, we observed a striking difference in the release of the reporter protein and gene, re-emphasizing the limitation of smaller size molecules (gadolinium) to fully represent the pharmacokinetics of larger size molecules (protein and gene). We observed significant improvement of biomarker release for both protein and ctDNA only for ML-CL group (1.5-fold for protein and 2.1-fold for ctDNA, **Fig. 7J-K**). This further underscores the importance of maximizing AE to improve biomarkers’ release – which is an effect that ML-CL can uniquely achieve while maintaining safety. It also confirmed that collection of larger blood volumes is critical for effective ctDNA detection. ctDNA degradation and clearance appear to be of lower significance, as long blood collection is close to sonication. In aggregate, these findings demonstrate that ML-CL, trained in mice, retains its safety and efficacy in tumor-bearing rats, hence revealing its translational potential for clinical diagnostic applications with FUS-enhanced liquid biopsy.

## Discussion

In this work, we developed and evaluated a data-driven closed-loop ultrasound nanotheranostic platform for transcranial interventions (**Fig. 1**). The proposed approach critically expanded the current state-of-the-art controllers (OL and CL) by integrating ML to markedly improve the treatment window and overcome rate-limiting barriers to the delivery of nanoparticles and detection of circulating-tumor DNA from the brain. Contrary to current practice ^75,76^, our findings indicate that the MB administration protocol is a major safety variable. Even under sonication settings that are generally considered safe under MB infusion, bolus administration can pose a significant risk, as the likelihood of broadband events can be substantially higher during the first pass of MB cloud – an effect that current simple (constant-pressure) clinical OL controllers cannot easily accommodate ^16,40,76,77^ unless they dramatically reduce the treatment window. Furthermore, OL controllers (constant-pressure) are prone to inconsistencies in harmonic emissions, including high variability (**Fig. 6E**) and a tendency towards overshooting (right-skewness; **Fig. 7E**), both of which are intensified by focal pressure uncertainties introduced by the skull. While in that respect CL controllers are more advantageous by adapting to real-time AE, they require careful target AE level selection ^16,17,39^ that makes them sensitive to cavitation threshold measurements. They are also bound to low target levels to avoid triggering broadband emissions (**Fig. 2I**). The proposed data-driven ML-CL platform overcomes these limitations. The ability to select target AE level, using a training dataset obtained from past sonications, and ability to predict broadband events makes the proposed data-driven closed-loop ultrasound platform not only immune to these fundamental tradeoffs in controlling MB dynamics but also able to scale to larger animal models. Together these advantages demonstrating the potential of data-drive feedback to be integrated into current clinical systems, provided there is sufficient data for training (e.g., current clinical systems already have these data ^37^) and, as such, can have an immediate and far-reaching impact on targeting CNS diseases.

Beyond the clinical implications of the proposed framework, by interpreting the trained MLP model (including additional harmonic components for broader perspective) it is possible to identify new or important features that shape MB dynamics, using model interpretation techniques such as SHAP approach. This can be used to enrich existing features for controlling the MB dynamics (e.g., ultra-harmonics and harmonics). It also revealed that the likelihood of inertial cavitation is greater when sonicating brain tumors, presumably due to higher MB accumulation or possibly due to abnormal vessel structure and mechanical properties. This finding indicates that calibrating the sonication settings in healthy brains and then applying them to diseased tissues can impact the MB dynamics in unexpected ways (inertial vs. stable cavitation) to undermine safety and confound mechanistic investigations that explore specific (immuno)mechanobiological interactions ^78^. It further underscores the importance of accounting for biological variability ^36,79^ and conducting systematic investigations for establishing treatment windows in disease models ^16^. SHAP analysis also revealed that pressure, while being a feature of high importance, its effect on the model’s output – hence, MB dynamics – is not monotonic (i.e., clear direction). That is, despite a weakly positive correlation, its effect on the model is likely to be influenced by complex interactions involving other MB AE features. Data-driven closed-loop ultrasound can help refine these findings and also support formal parametric analysis aimed at optimizing the FUS pulse sequences (excitation frequency, pulse duration, pulse repetition frequency) and MB properties for promoting desirable bioeffects ^60,80,81^. Collectively, our study demonstrates that machine learning models and data-driven analysis can form the basis for studying and analyzing the highly nonlinear and complex nanoscale microbubble dynamics in brain vessels, in addition to allowing for effective control of this highly nonlinear physical phenomenon.

Our investigation also demonstrated that the marked improvement in the therapeutic window of MB-FUS can have important implications for a wide range of therapeutic and diagnostic interventions in the brain. Most notably, as the ability to safely deliver a range of NP sizes (36 – 120 nm) can create new opportunities for targeting brain diseases and potentiate systemically administered novel gene editing nano-systems that incorporate larger and more potent cargos, thereby allowing them to effectively target a broad range of CNS diseases ^21^. Our analysis, which supported prior theoretical prediction that indicated higher delivery for 46 nm NPs under MB-FUS (**Fig. 4G**) ^82^, can be further explored to study other NP variables (e.g., surface charge, coating, loaded molecule(s)) that are critical for effective delivery in the brain. Moreover, considering that the AE levels were able to track the delivery of NPs across different species, can potentially alleviate the need for MRI verification and its significant logistic burden to the treatment. Thus, AI-powered ultrasound systems may support daily/weekly treatments, potentially at an outpatient and/or in limited resource settings, without compromising performance, thereby allowing the rapid and broad dissemination (i.e., similar to US imaging) and effective clinical translation of this transformative technology ^16,28,47,48^.

The level of robustness and flexibility offered by the proposed AI-powered ultrasound system can potentially be even more important for supporting FUS-enhanced liquid biopsy, where safety and portability can be crucial ^28^. Our study also revealed some important principles for the integration of FUS with liquid biopsy. Most notably, it clearly demonstrated the critical role of BBB permeability in restricting the release of cancer-soluble biomarkers. It also highlighted that protein based biomarkers (i.e., GLuc protein) that are characterized by low molecular size (∼20 kDa) ^83^ and high biomarker release per cell ^84^ are easier to detect. Unfortunately, GLuc DNA release was limited in mice. Several factors may have contributed to the limited detection of ctDNA in mice, including i) the larger size of this molecule (∼60kDa) ^71,72^, ii) the low number of copies (2 copy-limit per cell) ^85^, and iii) challenges associated with sample collection and assay sensitivity ^72,86,87^. However, as we showed in rats i) maximizing treatment window, ii) improving the assay sensitivity, and iii) increasing the blood collection volume (6.25x larger) are essential for improving ctDNA detection without compromising safety. Collectively, AI-powered ultrasound opens opportunities for repeated sono-interrogation of brain diseases using ctDNA and may provide a simple and economical way to confirm BBB opening independent from contrast-enhanced MR imaging. The improved performance observed at larger animals also suggests that scaling-up to humans can potentially lead to better responses, thereby creating further opportunities to genotype brain tumors and monitor the biology of therapeutic responses.

## Methods

### In vivo experiments

All animal procedures were performed according to the guidelines of the Public Health Policy on the Humane Care of Laboratory Animals and approved by the Institutional Animal Care and Use of Committee of Georgia Institute of Technology. 8-12 weeks old female C57BL/6J mice (Jackson Laboratory) were used in this study (N = 37 mice in total, and 114 mice in training dataset).

### USgFUS system

The system used in this study is a custom-built portable system that operates in two different modes: imaging mode and treatment mode.

In imaging mode, the system creates a 2D ultrasonic image by raster scanning (30 mm x 30 mm window) using the US imaging probe (Imasonic) that is coaxially aligned with the FUS transducer. To guide the FUS transducer to a target location in the brain, we overlay the 2D ultrasonic image onto MRI image using the eyes as a landmark. Desired target coordinates are determined relative to the center of the line connecting two eyes. After the relocation of the FUS transducer to such target coordinates, a pulse/echo scheme is used to align the ultrasound focus to the desired depth in the brain (relative to the skull) by determining the ultrasound travel time to the skull with sound speed of 1540 m/s. The targeting accuracy of this system is ± 500µm.

In the treatment mode of the USgFUS system, the imaging probe operates in passive mode and serves as a passive cavitation detector during BBB opening. When in passive mode, the recorded signal (11ms long) by the transducer is high pass filtered (cutoff 0.6 MHz) before it is fed to data acquisition system (Model 5000D, Pico Technology) and analyzed using FFT from the host computer. The FUS transducer (Sonic Concept) is driven by sinusoidal signal (0.5 MHz f_0_; 10 ms pulse length; 1 Hz PRF) generated by a function generator (Picoscope, Pico Technology), which has been amplified by a 50 dB power amplifier (Model 240L, Electronics & Innovation Ltd). The peak negative focal pressure of the FUS transducer in the water (free field) was determined using a calibrated hydrophone (2mm Model, Precision Acoustics).

The frequency bin we used for measuring broadband was 3.61 ± 0.3 KHz (7.22 f_0_), as this was closest to the maximum sensitivity of the PCD (3.5 MHz). The frequency range for ultra-harmonic and harmonic were determined as 0.5n multiples to f_0_ ± 0.3 KHz (where n is 3,4,5, …).

### Design of closed-loop AE controller for BBB opening

To attain a consistent level of MB dynamics, we incorporated a closed-loop controller that we have developed and validated in our previous studies ^17^. The controller has 3 main functions; it 1) mitigates for local AE fluctuation, 2) aims to achieve a target AE level (L^k^_target_; k depicts k^th^ harmonic) of a predefined observer state (k=7 in this study, which we selected from ML feature reduction), and 3) mitigates for global AE decay (MB kinetics).

The controller is also able to track cerebrovascular MB kinetics using the MB tracking pulse, P_tracker_. MB kinetics tracking allows an effective time window for controller’s operation. To prevent controller’s divergence caused by cerebrovascular MB clearance, we employed a MB tracking algorithm that monitors the MB kinetics after they are injected into the animals. The MB tracking pulse is a constant, low-pressure pulse (0.5 MHz; 10 ms pulse length; 1 Hz PRF; 60 kPa peak negative pressure) that follows the preceding controlled pressure pulse. The pressure for this pulse was chosen by selecting a pressure that resulted in minimal effect on BBB permeability, which was assessed using contrast-enhanced MRI. We have established criteria for controller’s operation: 1) when this MB tracking pulse detects 10 dB rise (relative to background signal) in 4^th^ harmonic (H4) level, controller is turned on; 2) after controller’s being turned on, initial slope of MB clearance is monitored for 40 s and is updated in real-time after the initial recording; 3) when the calculated slope indicates 20% decay from normalized maximum H4 level, the controller is ceased, and therapeutic pulse is maintained at its latest calculated pressure until the end of the sonication (130 Sec). For consistency, this P_tracker_ was also incorporated in all OL controllers in this study that started the controller operation with 10 dB rise in 4^th^ harmonic level.

### Machine Learning model training and testing

To establish a machine learning model for broadband emission prediction, a training dataset was formed using past AE data that utilized constant pressure sonication from total of 114 mice (10 ms pulse length, 1 Hz pulse repetition frequency, 0.5 MHz sonication frequency, 130s sonication). In order to find the relationship between current timepoint (t_n_) sonication to future timepoint (t_n+1_) broadband emission, features (i.e., input to machine learning model) were initially selected to be the current (t_n_) 1∼7^th^ ultra-harmonic (1.5∼7.5f_0_) emission levels, target region in the brain (x and y coordinates referenced to the eyes), pulse number, MB kinetics, pressure, and presence or absence of tumor (**Table 1**) – 2∼8^th^ (2∼8f_0_) harmonic emission levels were included for Shapley value (SHAP) analysis. Consequently, a label (i.e., ground truth for supervised classification machine learning) was selected to be binary presence of broadband emission (6 dB above baseline) at the next timepoint’s sonication (t_n+1_). The training dataset was then formed as N×D matrix, where N is the number of total AE dataset (N = 54,040), and D is our feature dimension (D = 20). The size of the corresponding label was (N×1). All AE levels were in logarithmic values. Our training dataset had a scarce presence of broadband emission above 6 dB, which was 8% (4,299/54,040) of the total dataset. To train the model under such scarcity, the model training dataset was under-sampled^55^, in which we randomly selected same number of AE data from the absence of broadband emission (label = 0) that matched 80% of the number of broadband emission presence (label = 1). To train MLP model, *feedforwardnet* and *train* functions in MATLAB (Mathworks, MA) were used with 10 neurons in 1 hidden layer (a simple proof-of-concept model), epochs size of 1000, and learning rate of 0.01 ^53^, using Lavenberg-Marquardt algorithm ^54^ to minimize mean squared error of the dataset. To test the trained model, the remaining 20% of dataset were used. Model testing was evaluated using a confusion matrix, specifically with accuracy (overall accuracy), sensitivity (i.e., recall, true positive vs. false negative), and precision (true positive vs. false positive), all of which are important metrics to evaluate machine learning models ^88^. Feature selection was performed using SHAP values with MATLAB *shapley* function ^56^, where we selected the top seven features of importance according to mean of SHAP values for feature reduction.

### Attentive Multilayer Perceptron (AMP)

Inspired by attention mechanisms, which have demonstrated the ability to selectively focus on the most relevant features of input data, leading to state-of-the-art performance across multiple domains, including natural language processing ^89^ and computer vision ^90^, we extended the MLP concept to the context-aware modeling. The proposed model integrates cross-attention to allow patient-specific features (query) to dynamically interact with frequency components (harmonics and ultra-harmonics) from broadband acoustic emissions (key-value pairs). This enables the model to selectively prioritize informative spectral biomarkers while suppressing noise, enhancing predictive accuracy for broadband emission and ensuring a controlled physiological state. The motivation for applying attention is that traditional MLPs treat all input features equally, whereas attention mechanisms facilitate dynamic feature interactions, allowing the model to adaptively weigh critical signals in relation to the individual characteristics. By leveraging cross-attention, this attention algorithm enables individual non-MBAE features (e.g., MB Kinetics, tumor presence, etc.) to be processed in the context of real-time environmental stimuli, allowing for an adaptive decision-making process to assess MB dynamics is at risk of instability.

The dataset was formed using past AE data that utilized constant pressure sonication from total of 114 mice. We categorize the input features into two groups: (1) Patient-specific features, which include the target region in the brain (x and y coordinates referenced to the eyes), pulse number, MB kinetics, pressure, and presence or absence of tumor; and (2) Real-time treatment feedback, which consists of broadband emission extracted from AE signals. These signals were selected to be the current (t_n_) 2∼8^th^ (2∼8f_0_) harmonic emission levels, 1∼7^th^ ultra-harmonic (1.5∼7.5f_0_) emission levels. The labels for supervised machine learning classification were binary indicators of broadband emissions exceeding 6 dB above the baseline at the subsequent sonication (t_n+1_). The training dataset was structured as an *N* x [*D*_l_, *D*_2_] matrix, where *N* presents the total number of AE samples (*N* = 54,040), *D*_l_ corresponds to the patient-specific feature dimension (*D*_l_ = 6), and *D*_2_ represents the real-time frequency-driven treatment feedback measurements (*D*_2_ = [7, 7]). The corresponding labels were stored as an *N* x 1 vector. Since broadband emissions above 6 dB were rare, accounting for only 8% of the total dataset (4,299 out of 54,040), we maintained this ratio when splitting the data into training (80%) and test (20%) sets to reflect the real data distribution. To address this class imbalance during training, we employed weighted cross-entropy loss, assigning a weight of 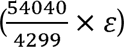 to broadband emissions exceeding 6 dB classes (in our case, we assign *ε* = 2). Additionally, we applied K= 10-fold cross-validation in training dataset to determine the optimal hyperparameters and select the best performance model. To mitigate overfitting, dropout, L2 regularization, and layer normalization were incorporated into the training process.

The AMP architecture is illustrated in **Fig. S3**. In this framework, individual-specific features first pass through a multilayer perceptron, encoding the information into a latent representation as the query (*Q*), while real-time frequency emission measurements are processed through another MLP, serving as the key (*K*) and value (*V*). The core of the architecture is the scaled dot-product mechanism, which enables interaction between these two feature sets. *Q* is derived from subject-specific features (*f*_l_), which encode the physiological state of the subject before sonication, *f*_l_ define the individualized treatment message that the model needs to consider when deciding whether MB dynamics are controlled or at risk of instability. *K* is from real-time frequency feedback (*f*_e_), reflecting the acoustic response from the AE measurements during the sonication, which contains harmonic and ultra-harmonic frequency measurement. The role of the *K* is to provide a reference against which the patient-specific information (*Q*) can be compared. The dot product computation of *QK^T^* produces a score matrix where each individual information interacts with each measurement in the latent space. After normalization via SoftMax, we can get attention scores that how much focus each individual feature should have on the corresponding real-time feedback. The *V* matrix contains the actual measured frequency emission information, and multiplying the attention score matrix by *V* produces an adaptive and feedback-aware feature representation for further classification. The key advantages of AMP lie in its ability to treat subject-specific and real-time feedback features separately while enabling context-aware interactions in the latent space. The attention mechanism employs three learnable transformation matrices, allowing the model to prioritize relevant frequency components dynamically, suppress noise, thus enhance feature selection automatically. Besides, the designed architecture ensures that the patient’s state is continuously re-evaluated in real time based on AE feedback, improving the accuracy and robustness of broadband emission prediction.

### Safety analysis of controllers

The performance of the ML-assisted closed-loop (ML-CL) controller was compared with 4 other controllers during MB-FUS BBB in healthy mice at two different target levels. The first groups were treated with state-of-the-art closed-loop (CL) controller ^17^. The second and the third group were open-loop (OL) controllers, and the exposure conditions were selected based on the average and maximum pressure used by ML-CL (P_Avg_ and P_Max_). The final group was also open-loop and used a pressure value from the cavitation threshold model (P_Model_) that corresponds to ML-CL target level. For fair comparison, all controllers (ML-CL, CL, P_Avg_, P_Max_, and P_Model_) had MB kinetics tracking function (constant 60 kPa pulse, 10 ms pulse length) that initiated the controller operation. As a safety feature, ML-CL reduces pressure after ML model’s prediction of broadband emission as well as after the occurrence (reaction) of broadband emission; CL and OL controllers reduced the pressure only after the occurrence of broadband emission. To compare the performance of the controllers 32 dB or 36 dB target levels were chosen from the cavitation threshold model (pressure vs. 7^th^ harmonic level). The first (32 dB) is expected to have a low level of broadband emission and the second (36 dB) is expected to have a high level. The key features of each controller are summarized in **Table 3**. For 32 dB operation, ML-CL (n=2), P_Avg_ (n=2), P_Max_ (n=2), and P_Model_ (n=2), totaling of 12 targets per group. For 36 dB operation, ML-CL (n=2), CL (n=2), P_Avg_ (n=2), P_Max_ (n=2), totaling of 8 targets per group. Animals from NP delivery were added to the group for K_trans_ analysis (6 targets per target level). For 36 dB operation in GFAP and Iba-1 staining, two targets were sonicated with ML-CL or P_Avg_ on right hemisphere and were compared with the contralateral (left hemisphere).

**Table 3-.** Different controller algorithms and the control law.

| Algorithm Name | Pressure | Control Law | Safety Feature |
| --- | --- | --- | --- |
| ML-CL | Varying | Closed-loop + MLP | Reaction + Prediction |
| CL | Varying | Closed-loop (Chapter 2&3) | Reaction |
| OL ($P_{Max}$) | ML-CL's Max Pressure ($P_{Max}$) | Reduce 5% pressure upon broadband emission | Reaction |
| OL ($P_{Avg}$) | ML-CL's Average Pressure ($P_{Avg}$) | Reduce 5% pressure upon broadband emission | Reaction |
| OL<br>($P_{\text{Model}}$) | Pressure from cavitation<br>threshold model<br>($P_{\text{Model}}$) | Reduce 5% pressure upon<br>broadband emission | Reaction |

### Experimental procedures

All sonications were performed with the FUS transducer operated at 0.5 MHz, with 10 ms pulse length and 1 Hz pulse repetition frequency, for 130 seconds under concurrent bolus intravenous (i.v.) administration of clinical grade lipid shell microbubbles (100 µL/kg, Definity, Lantheus Medical Imaging). For rat experiments, lab-made Definity-like MBs were used due to limited product supply. MB type, dose, and administration protocols were consistent with those used to collect the machine learning training datasets. During the sonications, we recorded/controlled the AE using the single element PCD. DCE-MRI was taken right after the sonication sessions. Mouse sonications: For 32 dB target level, six target regions in the brain (three in each hemisphere) were sonicated. For 36 dB target level, four target regions in the brain (two in each hemisphere) were sonicated. For NP delivery, four target regions were sonicated (32 dB on left and 36 dB on right). For brain tumor liquid biopsy, we performed five sonications (XY directions, separated by 1 mm) to cover the entire tumor and its periphery, and one healthy contralateral target was sonicated for targeting confirmation. Rat sonications: For NP delivery in tumors and brain tumor liquid biopsy, seven target regions were sonicated (XY directions separated by 1mm) to cover the entire tumor and its periphery.

### Dynamic contrast-enhanced magnetic resonance imaging (DCE-MRI)

To access the vessel permeability in the brain, we measured the volume transfer constant, K^trans^, by performing DCE-MRI (*Pharmascan 7T, Bruker,* IR, echo time, 2.5 ms; rep time, 1019.6 ms; flip angle, 30; FOV, 40 x 40 mm^2^) after the last sonication. Prior to injecting the contrast agents, we acquired background image in 6 different flip angles (2, 5, 10, 15, 20, and 30 degrees) to quantify T1 relaxation map, which is used for K_trans_ quantification. Consequently, we started collecting DCE-MRI with concurrent bolus administration of 8 μl gadolinium contrast agent (0.4 mL/kg, Prohance). The collected DCE-MRIs datasets were analyzed and K^trans^ values were calculated in Horos, using DCE tool plugin (Kyung Sung, Los Angeles, California). The arterial input function (AIF) was obtained based on Fritz-Hansen et al. method, as provided in the plugin ^91^.

### Brain tissue processing

Following MB-FUS treatments/sonications, the mice and rats were euthanized 2- or 6-hours post-treatment and transcardially perfused with 20 mL and 160mL of saline, respectively. For brain staining, perfusion continued with 4% paraformaldehyde (PFA) before harvesting the brains. Then, the brains were fixed with 4% PFA overnight at 4°C, followed by immersion in a 30% sucrose solution at 4°C until they sank to the bottom of the container. The brains were then embedded in O.C.T. compound and rapidly frozen to −80°C. Subsequently, 20 µm sections were cut using a cryostat (Leica 3050 S Cryostat) for further analysis. For NP delivery, brains were harvested after transcardial perfusion with saline and analyzed immediately (no exposure to PFA).

### Immunofluorescence / Immunohistochemistry staining and microscope imaging

To assess the safety and the biological effects induced by MB-FUS, immunofluorescence staining was performed on brain tissue slides. Tissues were first fixed in 4% paraformaldehyde at room temperature for 10 minutes. For sections requiring staining of intracellular markers (e.g., GFAP and Iba-1, which are markers for astrocyte and microglial activation for safety assessment), they were subsequently permeabilized with 0.1% Triton X-100 in PBS for 5 minutes. After washing with PBS, the sections were blocked for 1 hour at room temperature in a solution containing 1% Bovine Serum Albumin and 5% goat serum in PBS.

The sections were then incubated with the primary antibody of interest, diluted in 1% Bovine Serum Albumin (1:100), for 12 hours at 4°C. Following primary antibody incubation, the sections were incubated with the secondary antibody, diluted in 1% Bovine Serum Albumin (1:250), for 1 hour at room temperature. To stain the cell nucleus, samples were incubated with DAPI diluted in PBS (1:1000) for 10 minutes after washing. Finally, the sections were rinsed with PBS to remove excess antibody, mounted with mounting medium, and covered with coverslips. Samples were cured with mounting medium for 24 hours in the dark at room temperature before imaging. The sections were imaged with a 20x objective using a fluorescence microscope (Eclipse Ti2, Nikon). The quantification of the fluorescence images was performed using ImageJ.

Hematoxylin-eosin staining was performed to examine tissue damage and safety. Twenty-micrometer thick frozen sections (Leica 3050 S Cryostat) were dehydrated beforehand and stained using a Leica Autostainer (ST5010). The sections were imaged with a 20x objective using a brightfield microscope (Eclipse Ti2, Nikon).

### Statistical analysis

Results are expressed as means ± standard deviation (or SEM if stated under caption). Statistical analyses were performed using MATLAB and GraphPad Prism 7.0. For statistical analysis of two independent groups, an unpaired Student’s t-test was applied. For statistical analysis of self-comparison groups, a paired Student’s t-test was applied. For grouped analyses, we performed a One-way ANOVA, multiple comparison with Bonferroni’s correction. p < 0.05 was considered statistically significant (*p<0.5, **p<0.01, ***p<0.001, and ****p<0.0001, ns = not significant). For variance tests, f-test were used, and Levene’s test was used for skewed distributions. Data distribution was assumed to be normal (unless described in the manuscript for skewed distributions), but this was not formally tested in all data. Tumor-bearing animals were inoculated with tumors and then randomized prior to allocation to experimental groups. For the remaining studies, randomization was not relevant, as these experiments were conducted on uniform biological materials, such as commercially sourced cell lines. The investigators were not blinded to group allocation during experiments and outcome analysis. Blinding was not feasible for both the *in vitro* and *in vivo* studies, as these experiments were conducted by individual investigators who were aware of the experimental groups. Additionally, blinding was not considered relevant, as the data analysis was performed quantitatively (e.g., bioanalyzer quantification, PCR, and digital-PCR) and did not involve subjective, qualitative assessments. Animals were randomly assigned to experimental groups, but the experiments were not randomized, and the investigators were not blinded to group allocation or outcome assessment. Animal sample exclusion was minimized and only applied either when critical procedures failed or data fell below or above the negative or positive controls, respectively (see animal exclusion criteria section).

### Sample Exclusion Criteria

One mouse was excluded from the 36dB LB group due to mistargeting of the tumor, confirmed by contrast-enhanced MR imaging and observed in abnormal fluctuation of biomarker levels in the blood. Protein samples from 1 ML-CL rat were excluded due to luminescence signal below the negative control, included in the same Bioanalyzer run. ctDNA samples from 1 Pavg rat were excluded as quantification surpassed the positive control included in the same dPCR run. ctDNA samples from 1 ML-CL rat were excluded due to human error during the purification step. Overall, additional animals were placed in each of these groups to maintain n≥5.

### Nanoparticle preparation

To formulate PEGylated polystyrene nanoparticles, 20, 40, and 100 nm carboxylate-modified FluoSpheres™ (PS-COOH, Ex/Em: 580/605 nm, Invitrogen) were covalently modified with amine-terminated 5 kDa MW polyethylene glycol (methoxy-PEG5k-amine, Creative PEGWorks) by EDC carbodiimide chemistry, following a protocol we previously described ^92–94^. Briefly, PS-COOH nanoparticles (2 mg) were mixed with methoxy-PEG5k-amine (4-5x equivalent to total COOH groups on surface of PS-COOH particles) in 100 mM borate buffer (pH 8.2), followed by addition of excess sulfo-NHS (∼24x), and EDC (∼8x). Particle suspensions were placed on a rotary incubator and the reaction was allowed to proceed for 4 h at 25°C. After the reaction, particles were purified by ultracentrifugation (Amicon Ultra-15 mL 100 kDa MW cut-off, Millipore Sigma) with sterile PBS (pH 7.4) as the buffer (3 washes total). Finally, the purified nanoparticles were resuspended in sterile PBS (1 ml) and stored at 4 °C for subsequent characterizations and use in *in vivo* studies.

The physicochemical characteristics of PEGylated polystyrene nanoparticles were measured in 15x diluted PBS (∼10 mM NaCl, pH 7.4) at 25°C. Hydrodynamic diameter and ζ-potential (surface charge) were determined by dynamic light scattering and laser Doppler anemometry, respectively, using a Zetasizer NanoZS (Malvern Instruments, Southborough, MA). Particle size measurement was performed at 25°C at a scattering angle of 173° and is reported as the number-average mean. The ζ-potential values were calculated using the Smoluchowski equation and is reported as the mean ζ-potential. Detailed characterization shown in **Table 4**.

**Table 4-.**
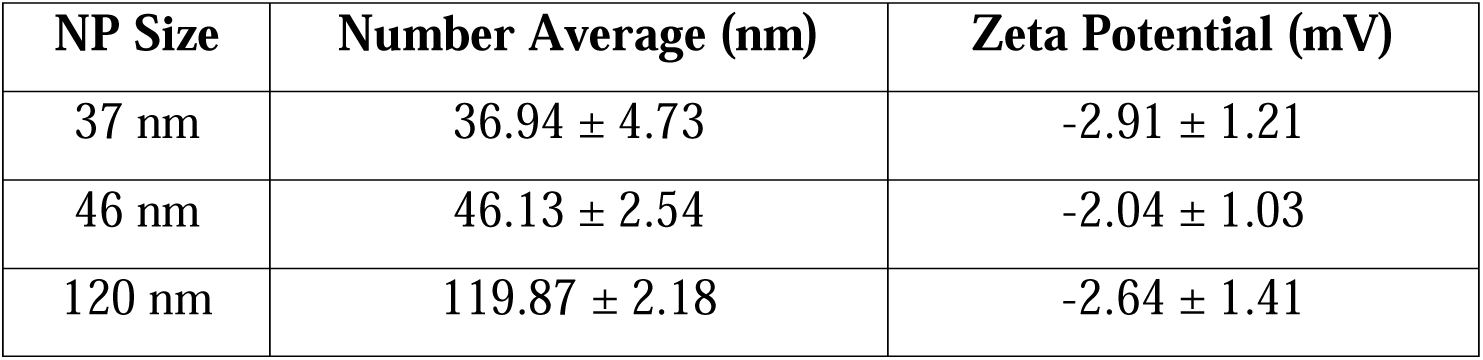
PEGylated polystyrene nanoparticle characterization.

| NP Size | Number Average (nm) | Zeta Potential (mV) |
| --- | --- | --- |
| 37 nm | 36.94 ± 4.73 | -2.91 ± 1.21 |
| 46 nm | 46.13 ± 2.54 | -2.04 ± 1.03 |
| 120 nm | 119.87 ± 2.18 | -2.64 ± 1.41 |

### Nanoparticle delivery and quantification

The sonications were performed as described previously, but the order of first sonicated side was alternated in between mice of the same group to account for MB accumulation. After the sonications were completed, each mouse received a 100µL intravenous bolus-injection of the designated NP size. PEGylated NPs were prepared for these injections at a concentration of 5 mg/kg. Considering an average body weight of 20g for mice and 160g for rats, each mouse received 0.1mg diluted in 100 µL and each rat received 0.8mg diluted in 800μL. The 37 nm, 46 nm, and 120 nm-NPs had stock concentrations of 1.3 mg/mL, 1.8 mg/mL, and 4.0 mg/mL. For mice, considering a 10% additional volume preparation, 84.61 µL of 37 nm-NPs were mixed with 25.39 µL of saline, 61.11 µL of 46 nm-NPs were mixed with 48.89 µL of saline, and 27.5 µL of 120 nm-NPs were mixed with 82.5 µL of saline, making a total of 110 µL of prepared NPs per injection. For rats, considering a 10% additional volume preparation, 676.9µL of 37 nm-NPs were mixed with 203.1µL of saline, 488.9µL of 46 nm-NPs were mixed with 391.1µL of saline, and 220µL of 120 nm-NPs were mixed with 660µL of saline, making a total of 880µL of prepared NPs per injection.Animals were euthanized 2h-post NP injection and transcardially perfused with 20mL (mice) or 160mL (rats) of saline.

To account for variations in size-dependent NP fluorescence signal, *in vitro* NP signal was assessed *a priori* for each size using the concentrations detailed above. Triplicate samples of 55µL of PBS and each NP size, same concentration from *in vivo* experiments, were individually placed into a black-walled, clear-bottom 96-well. Then, the well plate was placed in the Perkin Elmer IVIS Spectrum CT (Waltham, MA, USA) for fluorescent imaging of the brain. The machine was programmed using the Perkin Elmer Living Image 4.7.4 to capture fluorescent images with excitation at 570 nm and emission at 620 nm, exposure time of 0.5 second, lamp at low level, medium binning, C field of view, subject height at 1.00 cm, chamber temperature at 37 °C, and the lamp temperature at −90 °C. The machine was initially programmed to match the fluorescent particle’s original excitation of 580 nm and emission of 605 nm, but it automatically defaulted to the values above due to the range of filters installed. Measurements of total radiant efficiency were taken by placing the uniform region of interest (ROI) at the center of each well and covered the entire surface area. The color scale range was set from 2×10^9^ to 5×10^10^ (p/s)/(µW/cm^2^). The *in vitro* samples were used to normalize the fluorescent signal from 0 to 1 (highest measurement). Once normalized, the mean ratio of each size group was calculated and used to correct the recorded *in vivo* levels, as described below.

The harvested brains were immediately placed in the Perkin Elmer IVIS Spectrum CT (Waltham, MA, USA) for fluorescent imaging. The machine was programmed as before with exposure time of 10.0 seconds. The fluorescent image overlapped with the photograph of the brain. Measurements of total radiant efficiency were taken by placing the uniform ROI at the center of each sonication target. The color scale range was set from 9×10^7^ to 2.5×10^8^ (p/s)/(µW/cm^2^). To compare the delivery of NPs, the *in vivo* data was divided by the mean of the normalized *in vitro* data of each NP size group.

### Cell lines and cell culturing

GL261 glioma (Caliper Life Sciences, Hopkinton, MA, USA) and F98npEGFRviii cells (ATCC, Manassas, Virginia, USA) were separately cultured in Dulbecco’s modified Eagle’s medium supplemented with 10% fetal bovine serum (FBS) and 1% penicillin-streptomycin at 37°C and 5% CO_2_.

### Gaussia Luciferase transduction of GL-261 and F98-EGFRviii cell lines

GL-261 and F98-EGFRviii cell lines were transduced with Gaussia Luciferase (GLuc) using polybrene and the lentivirus cytomegalovirus humanized-GLuc (LV-CMV-GLuc-Puro, SignaGen Laboratories, Frederick, MD, USA) with the multiplicity of infection (MOI) equal to 2. Transduced cells were selected with overnight puromycin exposure at 10mg/mL.

### Tumor inoculation and growth in mice

GL-261-GLuc (mice) and F98-GLuc (rat) cells (10^5^ cells) were stereotactically implanted into the brain at 1-mm anterior, 1-mm to the right of the bregma, and 2-mm deep to skull bottom of 10-week-old female C57BL/6J mice (The Jackson Laboratory), or SRG (Sprague Dawley-Rag2 Il2rg/HblCrl)/Fisher CDF (F344/DuCrl) rats (Charles River). All tumors were allowed to grow up to 2.5 mm and then mice were evenly distributed among groups according to tumor volume that was quantified using the ellipsoid volume equation.

### Liquid Biopsy experiments

To assess the diagnostic potential of the proposed ML-CL controller, we employed it on GL-261-GLuc tumor-bearing mice (n = 6) and F98-GLuc tumor-bearing rats (n=7). Each animal was treated with 5 (mouse) or 7 (rat) sonication targets using ML-assisted controller to cover the tumor region at target levels established to be safe in healthy brains. To compare the role of FUS in brain tumor liquid biopsy, we split animals from the same cohort into a control group (n = 4 for mouse and 5 for rats), where no FUS was employed. These animals received the same MB injections with time intervals in between to represent each target sonication duration.

### Blood collection and Serum Isolation

For mouse, blood samples (225µL/each) were collected retro-orbitally 5 minutes before and 15 minutes after sonication of the treatment midpoint (for both FUS and control groups), and 2-hours-post-treatment (terminal – for FUS group) using EDTA-coated capillary tubes attached to a non-coated 1.5mL microcentrifuge tube. For rats, blood samples (1.2mL/each) were collected from the subclavian vein 5 minutes before and after sonication the treatment midpoint (for both FUS and control groups), using a 23G-needle attached to a luer slip syringe, and transferred to a non-coated 1.5mL microcentrifuge tube. All samples were allowed to coagulate in ice for 10 minutes prior to 1,000g centrifugation for 20 minutes. Serum was allocated for protein quantification (20µL – all animals) and ctDNA (80µL for mice and 500 µL for rats) purification/quantification.

### Protein Quantification

Serum (20µL) of each sample was placed in a single well of a black-walled, clear-bottom 96-well plate. Coelenterazine solution from the Pierce Gaussia Luciferase Glow assay kit (ThermoFisher, Waltham, MA, USA) was added to each well immediately prior to measurements using Synergy HT Microplate reader (Biotek Instruments, Santa Clara, CA, USA). Luminescence measurements were taken for 10 minutes at 30s intervals, with no lid, 1.0 mm height, 100ms delay, 135dB gain, and hole emission. Measurements for the first 5 minutes were considered and integrated, remaining measurements were disregarded due to signal decay after this timepoint.

### Circulating-tumor DNA (ctDNA) Purification and dPCR Quantification

Serum samples were placed in individual sterile 1.5mL microcentrifuge tubes, purified using the Plasma/Serum cell-free DNA Purification kit (Norgen Biotek, Ontario, Canada), and eluted to a final volume of 30µL. All samples were frozen overnight at −20°C.

Rat samples were pre-amplified using the Q5 Reaction kit (New England Biolabs, Ipswich, MA, USA) and forward and reverse primers, eluted in 10µM, with sequences 5’-AGAGATGGAAGCCAATGCCC-3’ and 5’-GTGCAGTCCACACACAGATC-3’, respectively. Each reaction included 10µL of the Q5 Reaction Buffer and 10µL of the Q5 High GC Enhancer (B9027S), 0.5µL of the Q5 High-Fidelity DNA Polymerase, 1µL of the Q5 dNTP Solution Mix, 0.5µL of each of the forward and reverse primers, 22.5µL of PCR-grade water, and 5µL of the template ctDNA sample. Reaction samples were placed in 0.5mL Eppendorf PCR tubes and run using Applied Biosystems’ Pro Flex PCR Thermocycler, which was programmed to process 1 cyle at 98°C for 30 seconds, and 10x cycles of 98°C for 10 seconds plus 68°C for 20 seconds plus 72°C for 15 seconds, and 1 cycle of 72°C for 2 minutes.

Custom fluorescent probe and sequence primers were synthesized for GLuc gene detection (Integrated DNA Technologies, Coralville, IA, USA). The probe, eluted at 100µM, has sequence of 5’-/56-FAM/CTGTCCCA/3IABIFQ-3’, and the forward and reverse primer sequences for GLuc gene, both eluted at 10µM, are 5’-ACCAGGGGCTGTCTGATCT-3’ and 5’– GGGATGAACTTCTTCATCTTGG–3’, respectively. Mouse and pre-amplified rat samples were processed for amplification and quantification. Each DNA sample was prepared by mixing 4µL of template ctDNA with 1.6µL of the probe, 0.32µL of forward and reverse primers, 10µL of QIAcuity dPCR Probe Master Mix (QiAgen, Hilden, Germany), and 23.76µL of PCR-grade water. All dPCR runs included *in vitro* positive control and negative control (PCR-grade water in place of DNA) samples. Samples were added to QIAcuity Nanoplate 26k 8-well/24-well plates and processed using QiAgen QIAcuity One digital-PCR (QiAgen, Hilden, Germany). dPCR was programmed to process FAM signal and included 1 cycle at 95°C for 2 minutes, and 40 cycles composed of 95°C for 15 seconds and 60°C for 30 seconds.

## Acknowledgments

We thank Dr. Johannes Leisen at Georgia Tech’s Magnetic Resonance Imaging core (MRI) facility for his outstanding technical support during FUS studies. We thank Dr. Andrew Kristof from the Immunological and Cellular Engineering Lab at Georgia Tech for his help with Gaussia Luciferase lentivirus transduction of F98-EGFRviii cell line.

## Author contributions

H.L., V.M., and C.A. conceived and designed the experiments; H.L., V.M. performed the experiments; H.L., V.M., C.K., S.Z., C.M.B., J.H.K., S.P., P.P. analyzed data; H.L., V.M., C.A. wrote the paper and prepared the figures; S.Z., and F.J.H. provided theoretical support; All authors reviewed and commented on the manuscript and approved the final version

## Data and materials availability

All data needed to evaluate the conclusions in the paper are present in the paper and/or the Supplementary Materials. Any additional requests for information can be directed to, and will be fulfilled by, the corresponding authors. Source data are provided with this paper.

