## Supplementary Materials for "Data-driven feedback augments ultrasound nanotheranostics in brain tumors"

**Materials and Methods**

1. **Training of different machine learning models with past dataset**

Past acoustic emission (AE) datasets from blood-brain barrier (BBB) opening experiments can be used to train different machine learning (ML) models. Apart from the multilayer perceptron (MLP) model ^1^ that we mainly incorporated in the study, we additionally trained support vector machine (SVM) ^2^ models in MATLAB (**Fig. S1**). All types of models showed similar accuracy, precision, recall (sensitivity), specificity, and f1 score (**Fig. S2**). They were also trained in 1) different kernels for SVM – RBF and linear, 2) different loss for MLP – cross entropy (MLP-CE) and mean squared error (MLP-MSE), and 3) different minority sampling methods, such as synthetic minority oversampling technique (SMOTE) and undersampling. Using SMOTE tend to have high precision (false positive vs. true positive), but lower sensitivity (true positive rate).

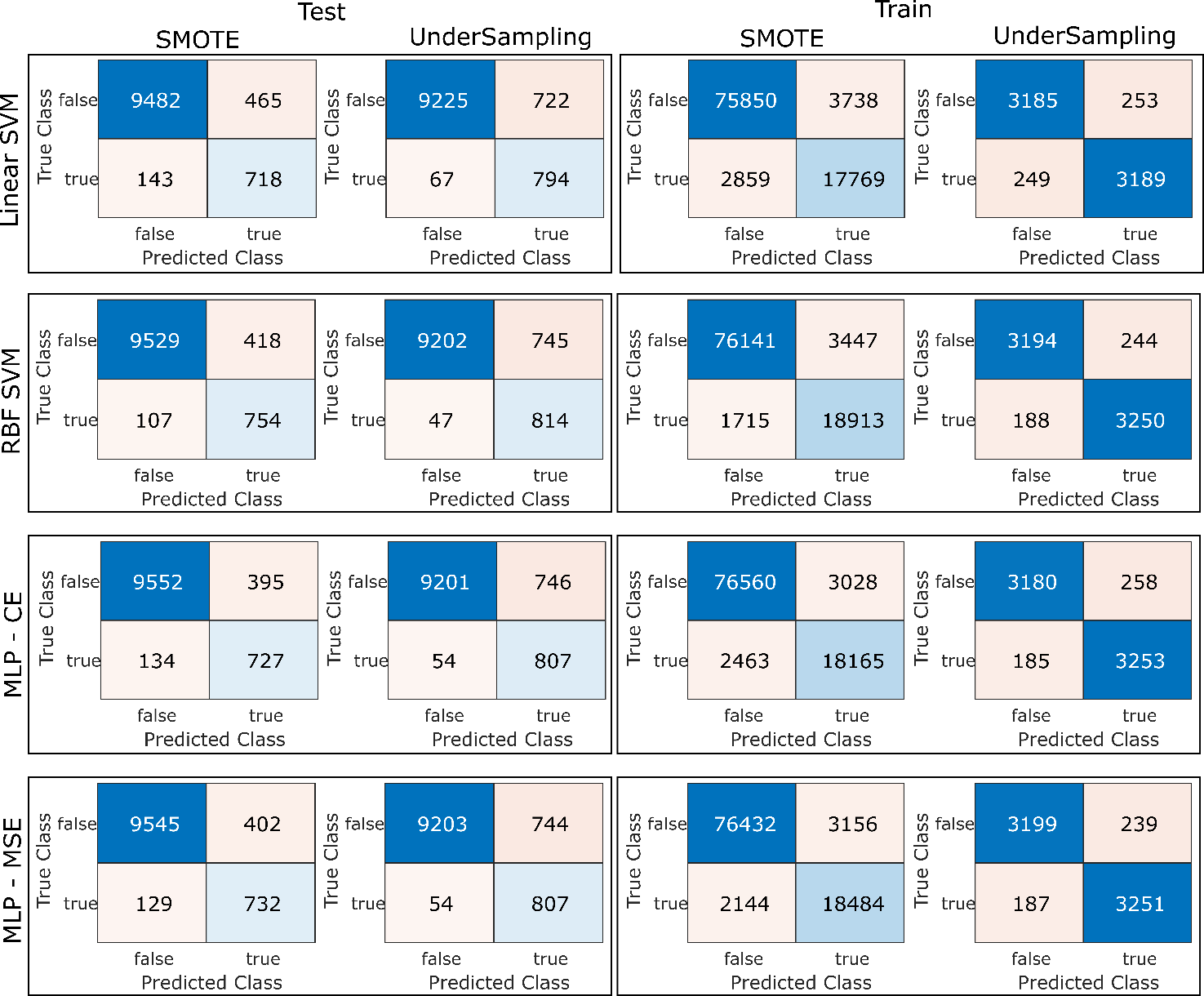

**Fig. S1.** Evaluation of different ML algorithms on training dataset: Linear kernel SVM, RBF kernel SVM, MLP with cross-entropy, and MLP with mean squared error (MLP-MSE, the model used for ML-CL). Left column shows performance on the testing dataset, and right column shows performance on the training dataset. For each algorithm, the effect of undersampling or synthetic minority oversampling (SMOTE) was also compared.

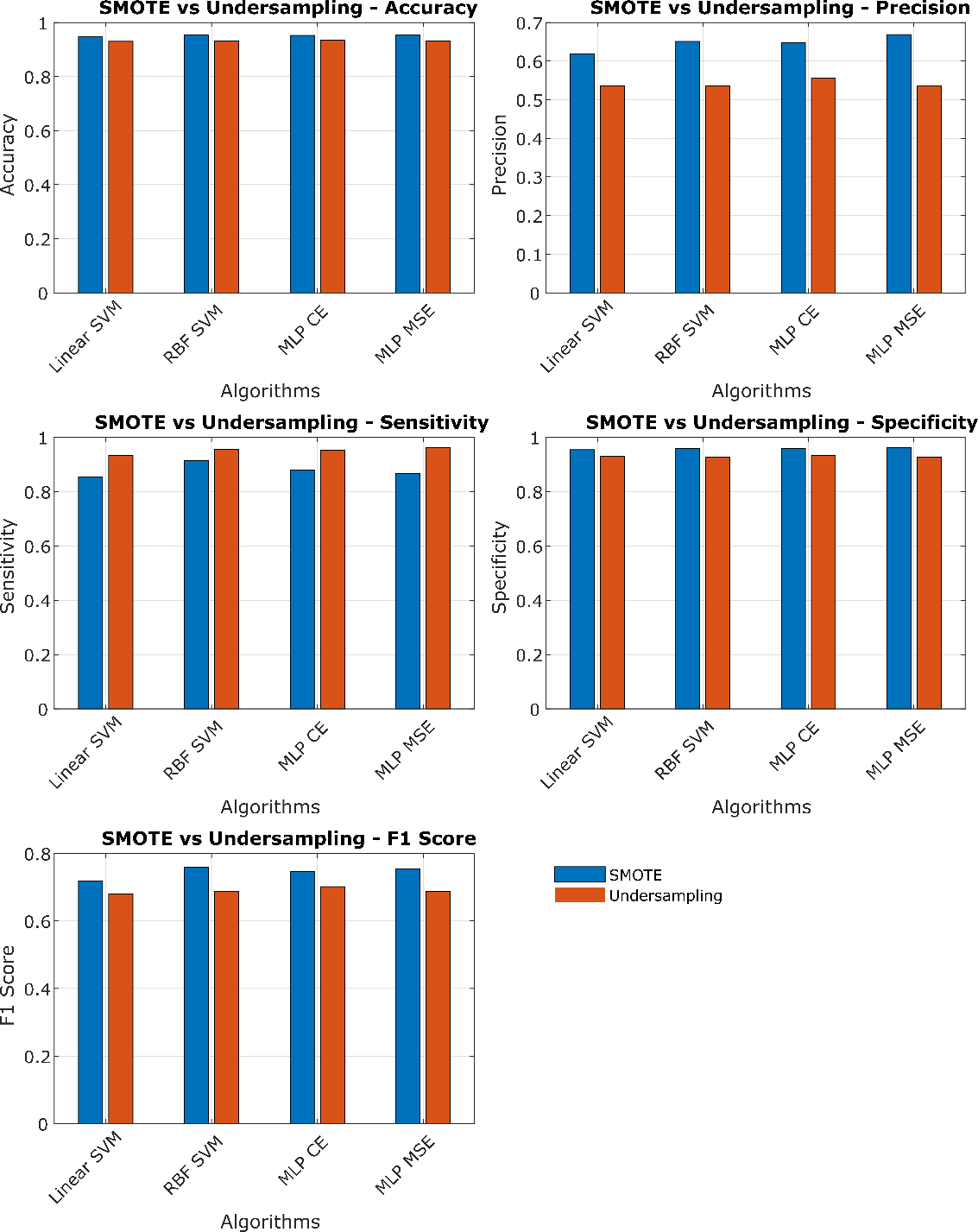

**Fig. S2.** Quantification of each ML algorithm's performance. All algorithms showed similar metrics: Accuracy, precision, specificity, sensitivity, and F1 score, given the same minority sampling method. Algorithms trained with SMOTE sampling technique showed tendency of low false positive, but high false negatives.

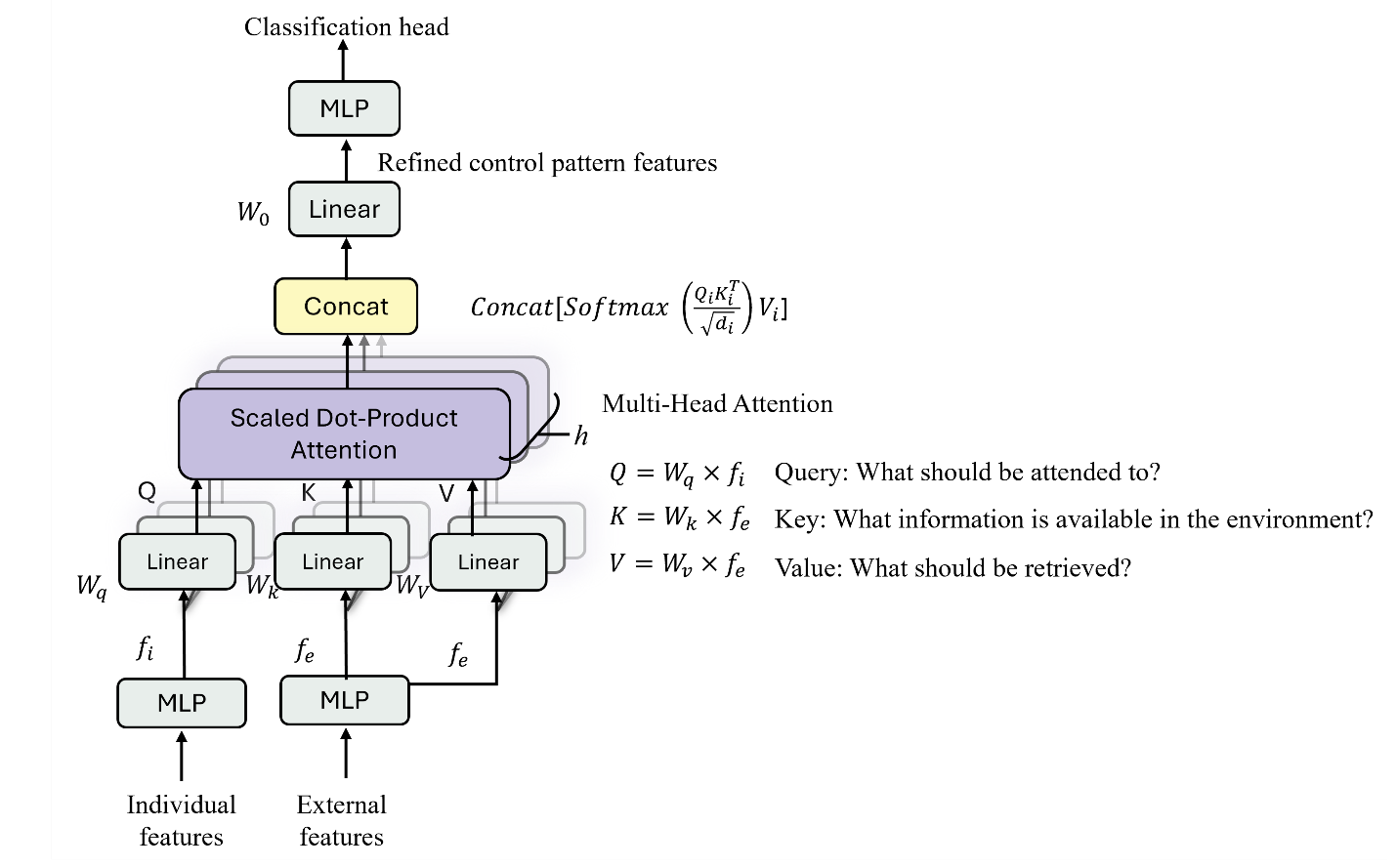

**Fig. S3.** Attention multilayer perceptron (AMP) architecture.

3. **Evaluation of attentive multilayer perceptron (AMP)**

The dataset was formed using past AE data that utilized constant pressure sonication from a total of 114 mice. We categorize the input features into two groups: (1) Patient-specific features, which include the target region in the brain (x and y coordinates referenced to the eyes), pulse number, MB kinetics, pressure, and presence or absence of tumor; and (2) Real-time treatment feedback, which consists of broadband emission extracted from AE signals. These signals were selected to be the current (t_n_) 2~8^th^ (2~8f_0_) harmonic emission levels 1~7^th^ ultra-harmonic (1.5~7.5f_0_) emission levels. The labels for supervised machine learning classification were binary indicators of broadband emissions exceeding 6 dB above the baseline at the subsequent sonication (t_n+1_). The training dataset was structured as an $N\times[D_{1}, D_{2}]$ matrix, where $N$ presents the total number of AE samples ($N=54,040$), $D_{1}$corresponds to the patient-specific feature dimension ($D_{1}=6$), and $D_{2}$ represents the real-time frequency-driven treatment feedback measurements ($D_{2}=[7, 7]$). The corresponding labels were stored as an $N\times1$ vector. Since broadband emissions above 6 dB were rare, accounting for only 8% of the total dataset (4,299 out of 54,040), we maintained this ratio when splitting the data into training (80%) and test (20%) sets to reflect the real data distribution. To address this class imbalance during training, we employed weighted cross-entropy loss, assigning a weight of ($\frac{54040}{4299}\times\varepsilon$) to broadband emissions exceeding 6 dB classes (in our case, we assign $\varepsilon=2$). Additionally, we applied K= 10-fold cross-validation in the training dataset to determine the optimal hyperparameters and select the best performance model. To mitigate overfitting, dropout, L2 regularization, and layer normalization were incorporated into the training process.

We found that compared to MLP, AMP had a decreased false positive rate (**Fig. S4A**). Moreover, SHAP analysis of the AMP model (**Fig. S4B**) provided similar results to our MLP SHAP analysis (**Fig. 1D**).

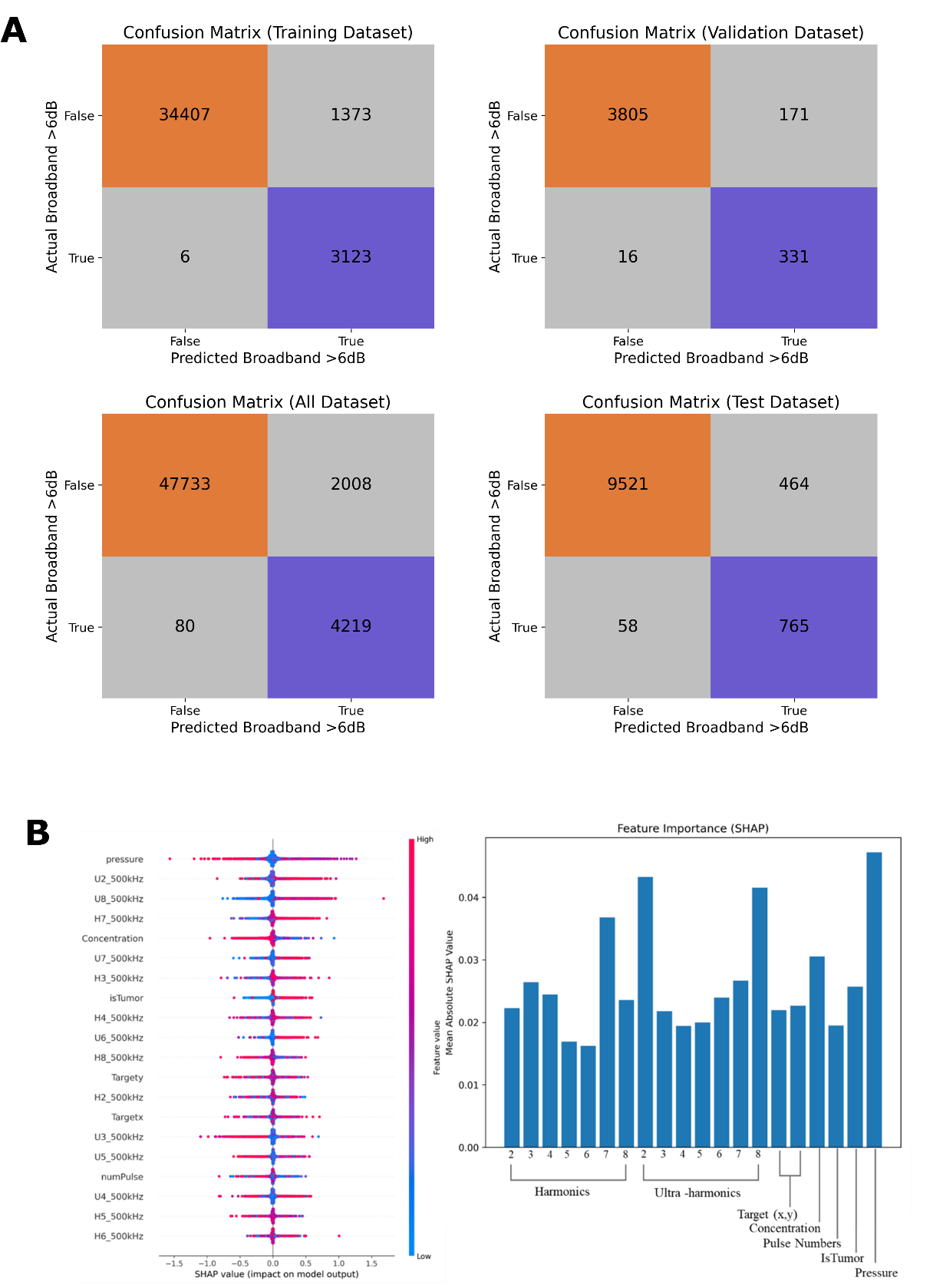

**Fig. S4.** A) Confusion matrix of AMP in training, validation, testing, and overall dataset. Compared to MLP only, AMP showed general improvement in lower false positives. B) Shapley additive analysis (SHAP) onto AMP. MLP and AMP had similar importance in top features.

4. **Correlation between MB kinetics and broadband emission strength obtained from the training dataset**

We analyzed the training dataset further to find the relationship between broadband emission strength and MB kinetics. We correlated the MB kinetics (transient decay in harmonic emission, normalized to maximum harmonic emission level) to broadband emission. Our analysis showed that stronger broadband emission (>15 dB) has maximum likelihood at higher MB kinetics (i.e., right after bolus injection of MBs) (**Fig. S5**). This may suggest the inherent risk of using constant pressure sonication; at the least, constant pressure sonication should be started after a few seconds of MB administration.

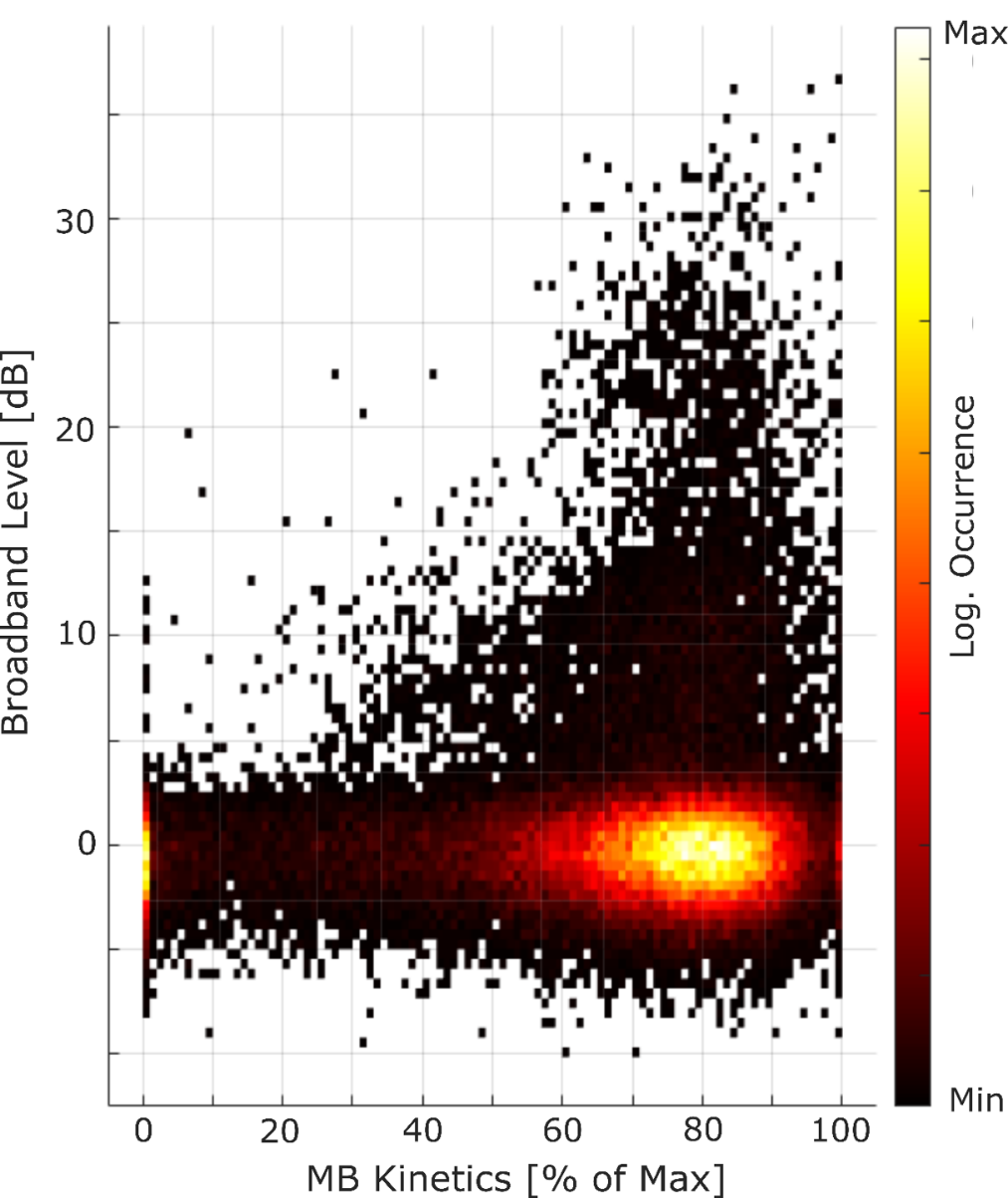

**Fig. S5.** 2D histogram of Broadband emission level as a function of MB kinetics. Colorbar indicates the logarithm of the number of occurrences.

5. **Application of machine learning-assisted closed-loop controller (ML-CL) onto healthy mouse with 32 dB target level**

To evaluate the operation of the ML-assisted closed-loop controller (ML-CL), we first performed MB-FUS mediated BBB opening in healthy mice by targeting three regions in each brain hemisphere using the USgFUS system that we used to collect the training data. We selected a target level of 32 dB at 7^th^ harmonic emission using the cavitation threshold curve identified from our training dataset (**Fig. S6A**). We chose this target level because it was at the highest level within the linear regime of the curve. With a 32 dB target level, we found that the average and maximum pressure decisions (P_Avg_ and P_Max_, as reported as peak negative pressures) of ML-CL were 0.17 MPa and 0.23 MPa, respectively (**Fig. S6B**). Along with the pressure corresponding to 32 dB in the cavitation threshold model (P_Model_ = 0.14 MPa, **Fig. S6A**), these pressures provided the necessary exposure conditions for implementing constant pressure open-loop controllers (OL) to compare and benchmark the ML-CL controller’s performance. This was performed by comparing the resulting AE, total broadband emission events, and BBB opening strength quantified using dynamic contrast-enhanced MRI (DCE-MRI) in healthy mice (**Fig. S6C-D**). Note that every controller incorporated MB kinetics tracking pulse (a small constant pressure pulse to monitor MB kinetics) to begin their operation upon detection of MB arrival to the brain (**Fig. S6B**).

We found that ML-CL achieved 31.6 ± 0.6 dB 7^th^ harmonic emission level with significantly lower AE fluctuation (i.e., more stability and ability to achieve and maintain prescribed level) compared to P_Model_ (24.4 ± 3.5 dB) and P_Avg_ (27.5 ± 4.7 dB) OL controllers (**Fig. S6E**), because of the feedback algorithm. In line with the AE levels, we observed a similar trend in K_trans_ values derived from DCE-MRI across different controllers (**Fig. S6F-G**). Although P_Max_ (31.7 ± 0.7 dB) showed comparable AE performance and higher K_trans_ compared to ML-CL, it also exhibited 3 instances of broadband emission during sonication (**Fig. S6H**), which indicates that safety was compromised. On the other hand, we observed that ML-CL controller had predicted and responded to 0.2% (3/1820) instances of potential broadband emission events (**Fig. S6I**) during its real-time operation. Post-sonications analysis further indicated that if it was integrated into the OL controller, the MLP model could have predicted all broadband emission events (3/4680) during the OL controller’s operation (**Fig. S6I**).

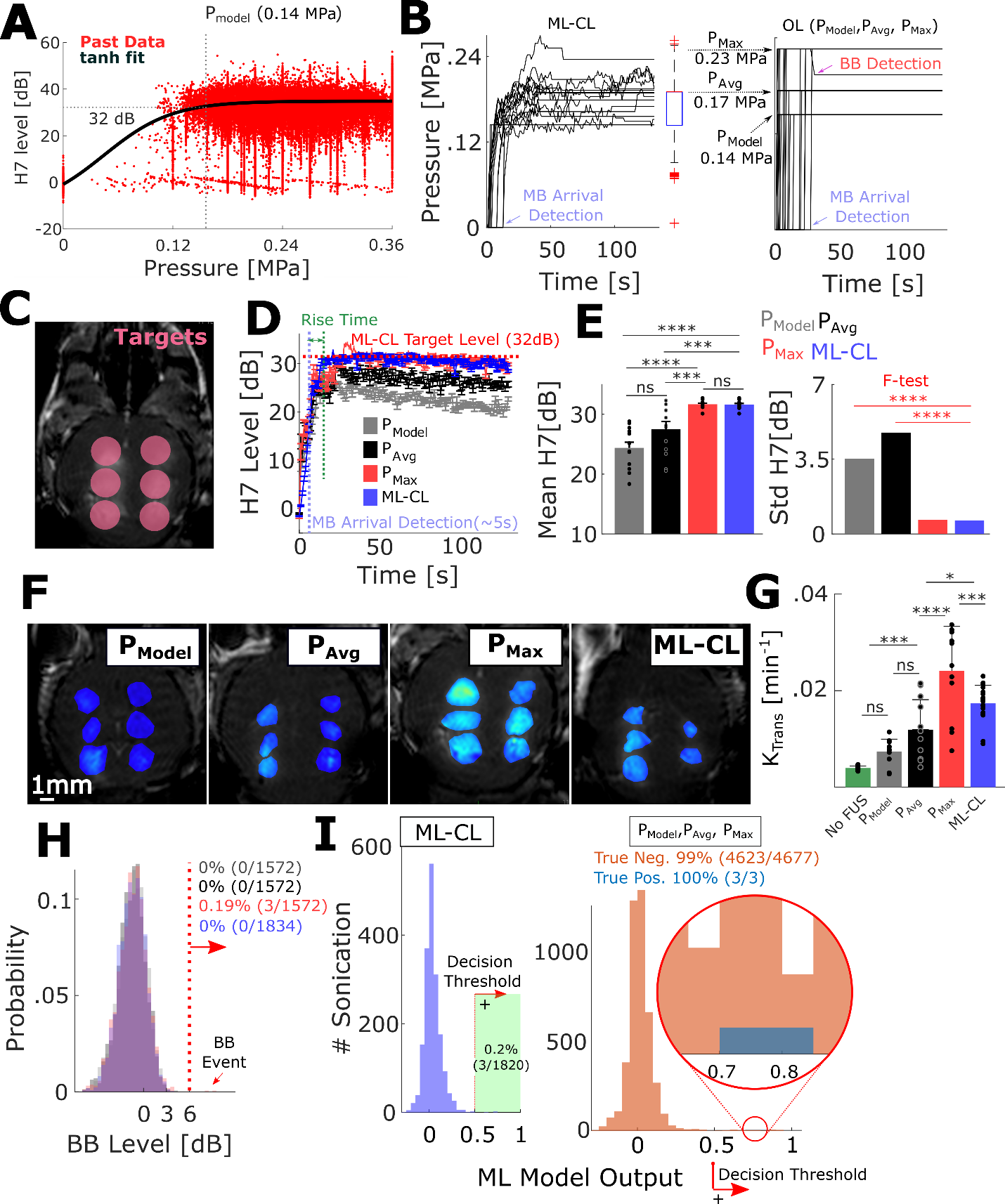

**Fig. S6.** Performance assessment of ML-CL in BBB opening with 32 dB target level. **A)** Cavitation threshold model from the training dataset. 7^th^ harmonic AE as a function of pressure. The model can be used to determine an adequate target level for the controller, and the P_Model_ is determined here as 0.14 MPa. **B)** ML-CL pressure decisions for 32 dB target level (left). The average and maximum pressures used by ML-CL are used to determine P_Max_ (0.17 MPa) and P_Avg_ (0.23 MPa) for OL controllers (right). A decrease in pressure in OL indicates a reaction to broadband emission event (> 6dB). **C)** Representative MRI image for 32 dB target level sonication targets. 3 targets in each brain hemisphere (total 6 targets per mouse) were treated. **D)** 7^th^ harmonic emissions during sonication for each controller. n = 2 animals (total 12 targets) per group. **E)** Mean (left) and standard deviation (right) of 7^th^ harmonic emission for ML-CL. A variance test (f-test) was performed to compare standard deviation (right). **F)** MRI T1 images using the controller at 32 dB or equivalent OL pressure. **G)** Quantification of K_trans_ values through DCE-MRI. **H)** Histogram for broadband emission levels during sonication. Broadband emissions higher than 6 dB were considered a broadband emission event, whose probability for each controller is highlighted next to the dotted line. **I)** Histogram for MLP decision during ML-CL sonication (left). A model output of 0.5 or greater was considered a positive prediction. For this target level (32 dB), the model predicted 0.2% (3/1820) of sonication. Application of MLP onto OL algorithms (right). Orange indicates true negative accuracy (predicting no broadband emission) and blue indicates true positive accuracy (predicting broadband emission). *p<0.05, **p<0.01, ***p<0.001, and ****p<0.0001. ns = not significant. Statistical analyses were performed through One-way ANOVA and Bonferroni correction.

6. **Post-sonication application of MLP onto 36 dB OL controllers**

We applied MLP to predict broadband emissions on the acquired AE dataset after OL (P_Max_ and P_Avg_, 0.33 and 0.25 MPa, respectively) sonication. MLP was able to predict 51% of the existing broadband emission events that were observed during sonication (**Fig. S7**).

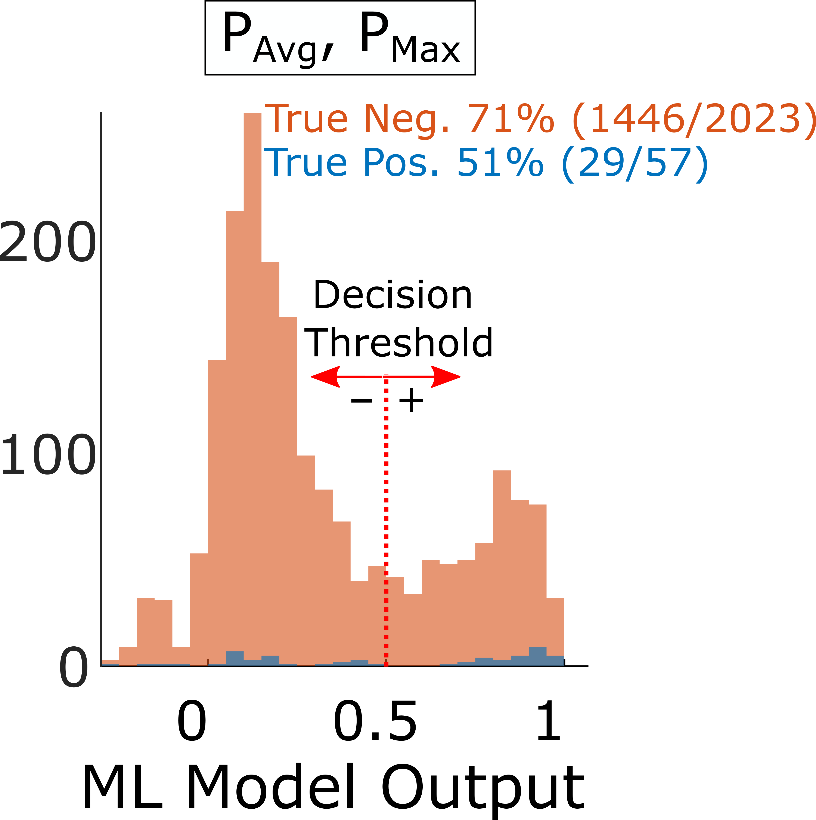

**Fig. S7.** Post-sonication application of MLP onto OL (P_Max_ and P_Avg_, 0.33 and 0.25 MPa, respectively) sonication. MLP was able to predict 51% of the existing broadband emission events that were observed during sonication. Orange indicates true negative accuracy (predicting no broadband emission), and blue indicates true positive accuracy (predicting broadband emission).

7. **Broadband emission and real-time pressure**

For more detailed information of **Fig. 4J**, pressure vs. broadband level throughout all sonication AE datasets (5240 in total) was included in this Supplementary Information with a more detailed figure of **Fig. 4J** with individual data points (**Fig. S8**). The majority of data (due to scarcity of broadband emissions) were around noise level (less than 4 dB). Thus, the broadband emissions for each controller at the noise level were downsampled with a 1% rate to obtain the fit of the hypertangent curve. There was 34% increase in the inflection point (i.e., curve shift), whose pressures were used to determine the upperbound of the therapeutic window.

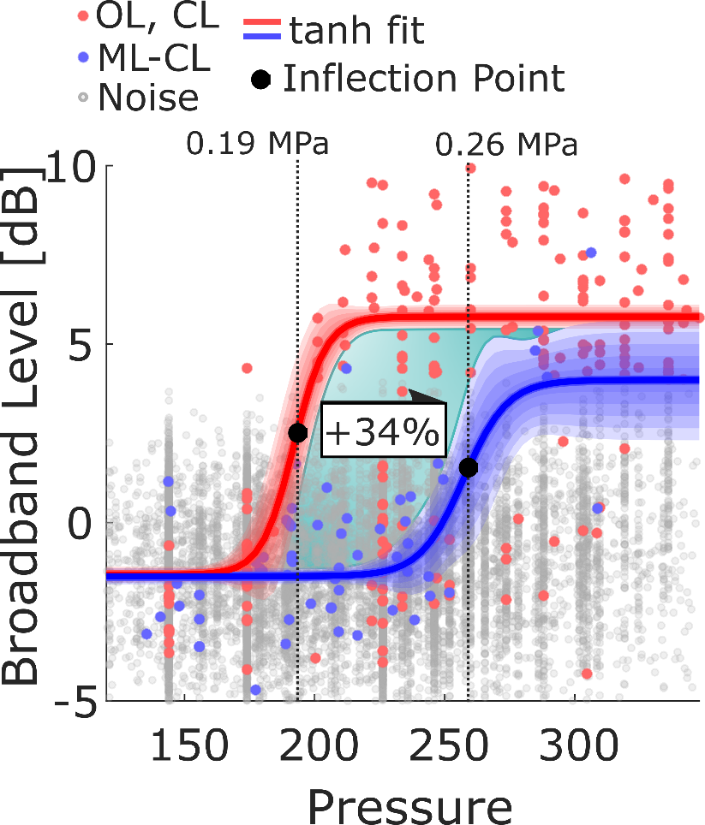

**Fig. S8.** Pressure vs. broadband level throughout all sonication AE dataset (5240 in total). A more detailed figure of **Fig. 4J** with individual data points. The majority of data (due to scarcity of broadband emissions) were around noise level (less than 4 dB). Thus, the broadband emissions for each controller at the noise level were downsampled with 1% rate to obtain the curve. There was 34% increase in the inflection point (i.e., curve shift), whose pressures were used to determine the upperbound of the therapeutic window.

8. **Performance of ML-CL in safety**

In assessing ML-CL’s safety with 36 dB, we assessed whether there were any false negatives (no broadband but tissue damage) after sonication. We found that CL contained 2 out of 8 targets where no broadband was observed but had hemorrhage (**Fig. S9**). This case was also found at P_Avg_ sonication.

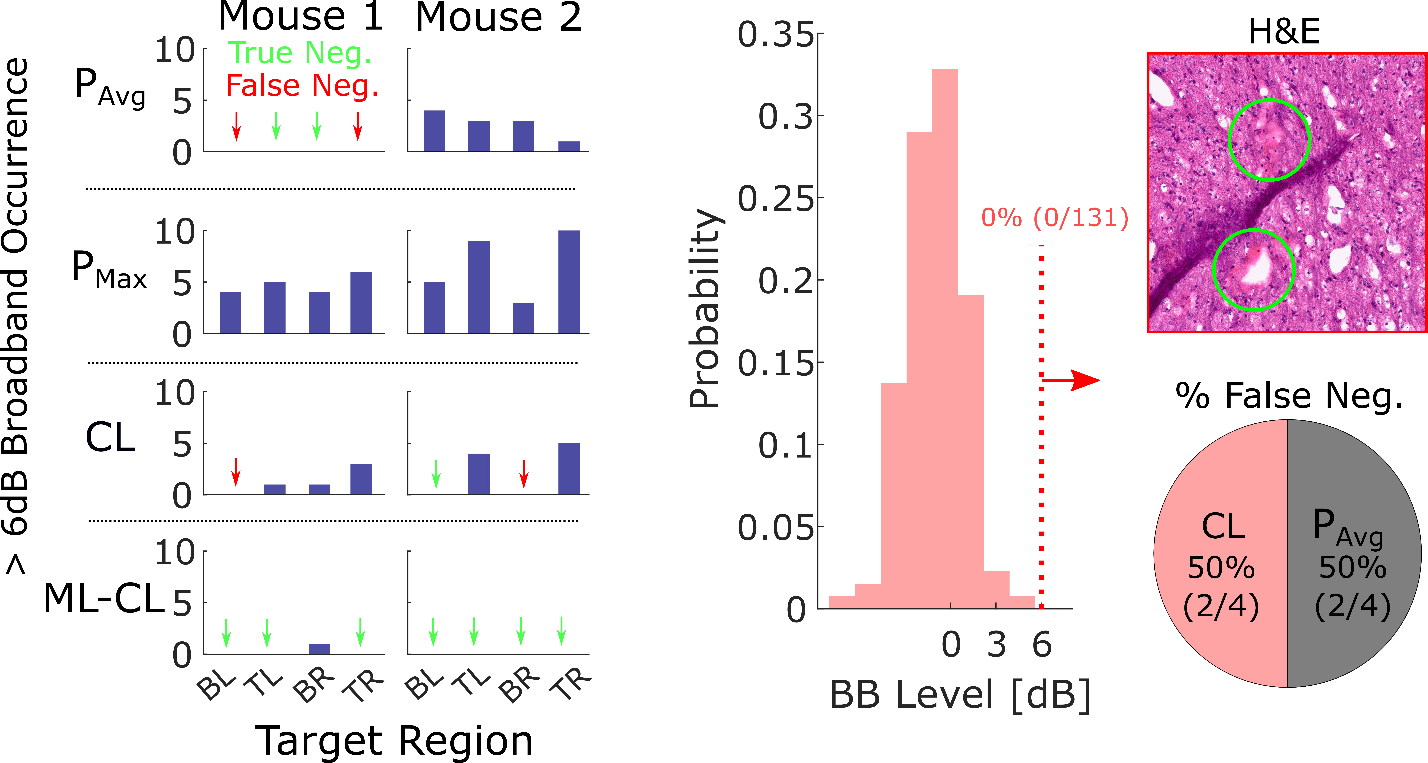

**Fig. S9.** Presence of broadband emission at BL (bottom-left), TL (top-left), BR (bottom-right), and TR (top-right) target locations. Green arrows indicate true negative (no broadband and no damage), and red arrows indicate false negative (no broadband and damage). Also, a Histogram of broadband emission for BL target in CL algorithm where no broadband was observed but petechiae was observed. Bottom right: pie chart for each controller’s stake in false-negative events.

9. **Performance of ML-CL in nanoparticle delivery**

In applying ML-CL onto different sizes of nanoparticle delivery, we applied 32 dB and 36 dB target level ML-CL onto the left and right side of healthy brain hemispheres, respectively. The pressure used by ML-CL for each the target levels was notebly different (**Fig. S10A**), which resulted in significantly different (p=0.009) 7^th^ harmonic emissions (**Fig. S10B**). Broadband emission event (>6dB) was absent except for one event at 120 nm sonication with 36 dB ML-CL (**Fig. S10C**).

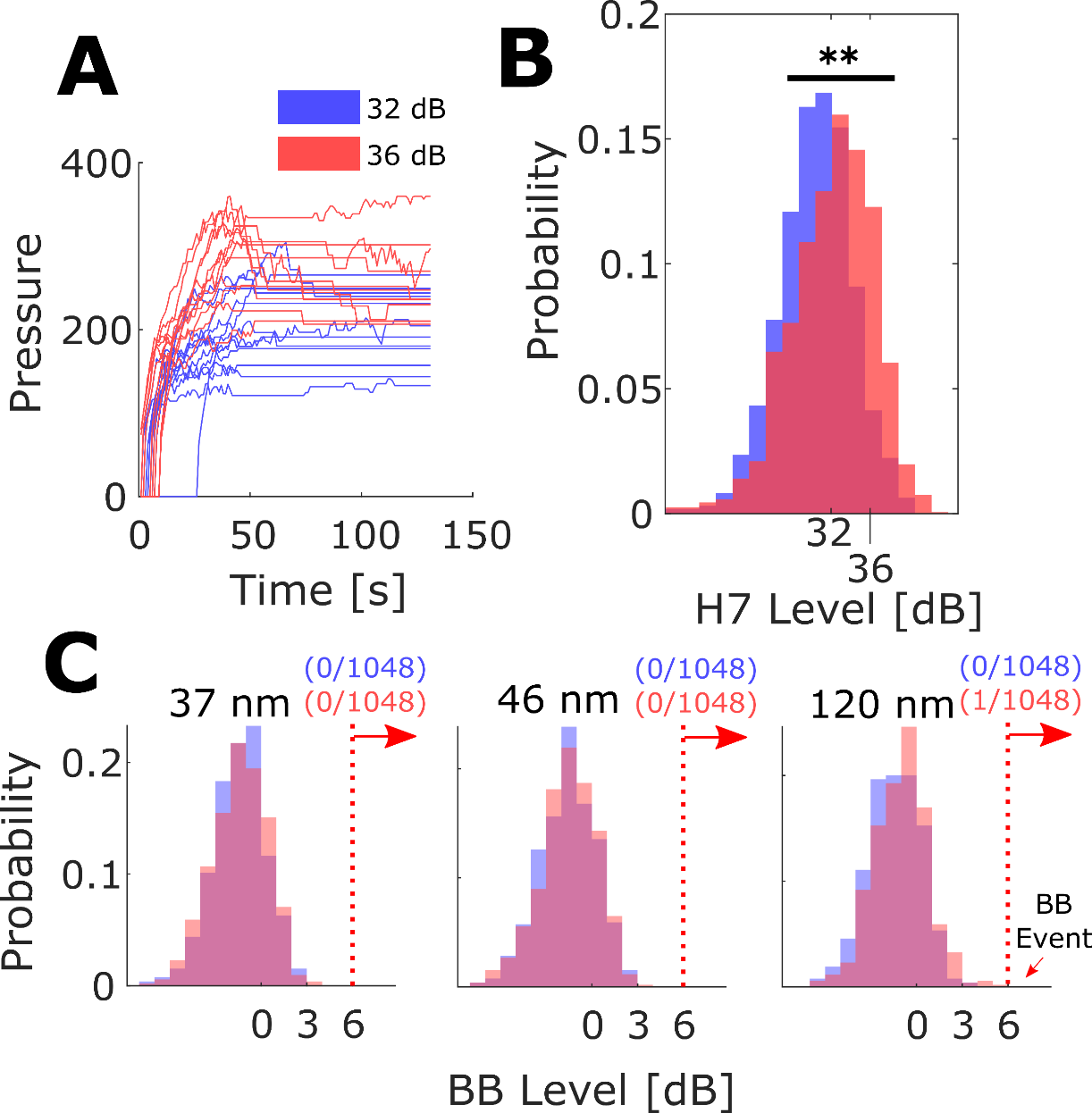

**Fig. S10. A)** Pressures used by ML-CL at each (32 and 36 db) target level. **B)** Histogram for 7^th^ harmonic emission during ML-CL sonication. **C)** Histogram for broadband emission during ML-CL sonication. One event was present at 120 nm cohort during 36 dB target level operation. All red colors indicate 36 dB target level, and blue colors indicate 32 dB target level.

10. **Liquid biopsy (LB) with multiple timepoint collection in mice**

For mice, blood samples (225µL/each) were collected retro-orbitally 5 minutes before and 15min after sonication at the treatment midpoint (for both FUS and control groups) and 2-hours-post-treatment (terminal – for ML-CL group) using EDTA-coated capillary tubes attached to a non-coated 1.5mL microcentrifuge tube. All samples were allowed to coagulate in ice for 10 minutes prior to 1,000g centrifugation for 20 minutes. Serum was allocated for protein quantification (20µL – all animals) and ctDNA (80µL for mice and 500 µL for rats) purification/quantification.

We found that the protein concentration in the blood increased by 3.1-fold immediately after sonication, as compared to pre-sonication (p<0.05) for 36 dB ML-CL (**Fig. S11**), indicating a burst release of GLuc protein from the tumors to the circulation following ML-CL sonication. Although these levels persisted even after 2 hours, a declining trend was evident (**Fig. S11**). In contrast to protein, the GLuc gene analysis showed that, on average, the levels in the number of positive droplets in the circulation with and without MB-FUS were very low and inconsistent.

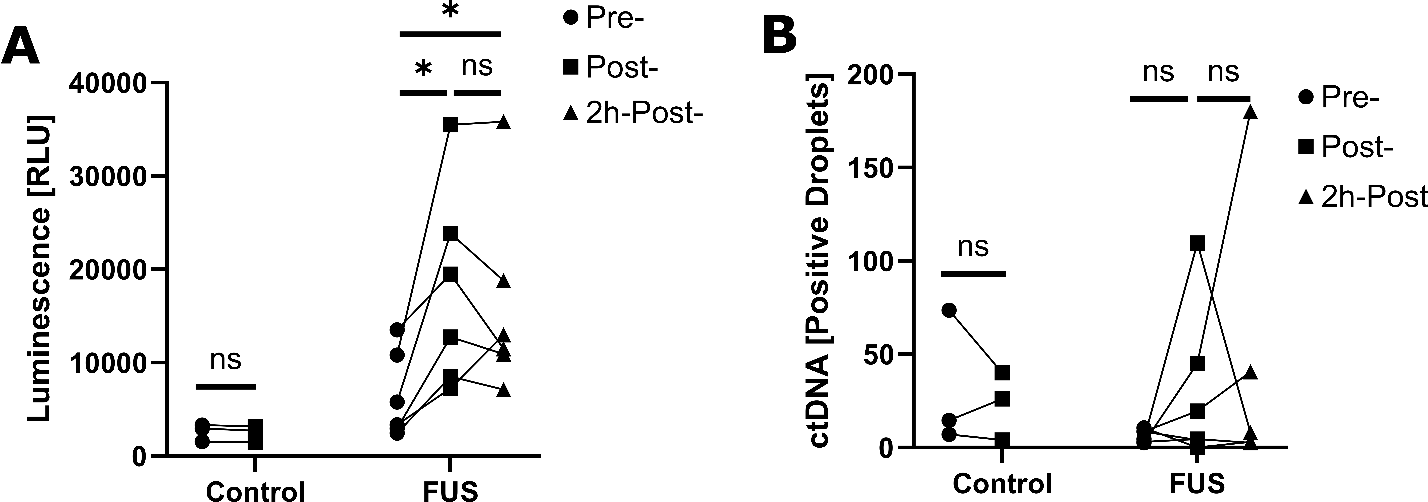

**Fig. S11.** Protein and ctDNA quantification at multiple timepoint. Control group includes pre- and post-treatment samples, while FUS group includes pre-, post-, and 2h-post **A)** Protein quantification with luminescent signal acquired (see methods for details of quantification). **B)** ctDNA quantification (positive dPCR droplets).

11. **Liquid biopsy (LB) rat strain impact of biomarker baseline concentration**

Biomarker quantification in different rat strains (immunocompetent and immunocompromised) indicated that the protein concentration baseline is elevated for immunocompetent rats (**Fig. S12**), which is probably due to the low clearance of molecules in circulation^5,6^ and retention of biomarkers. The saturation of biomarker concentration does not represent the realistic application of this technique and may diminish the impact of FUS on biomarker release.

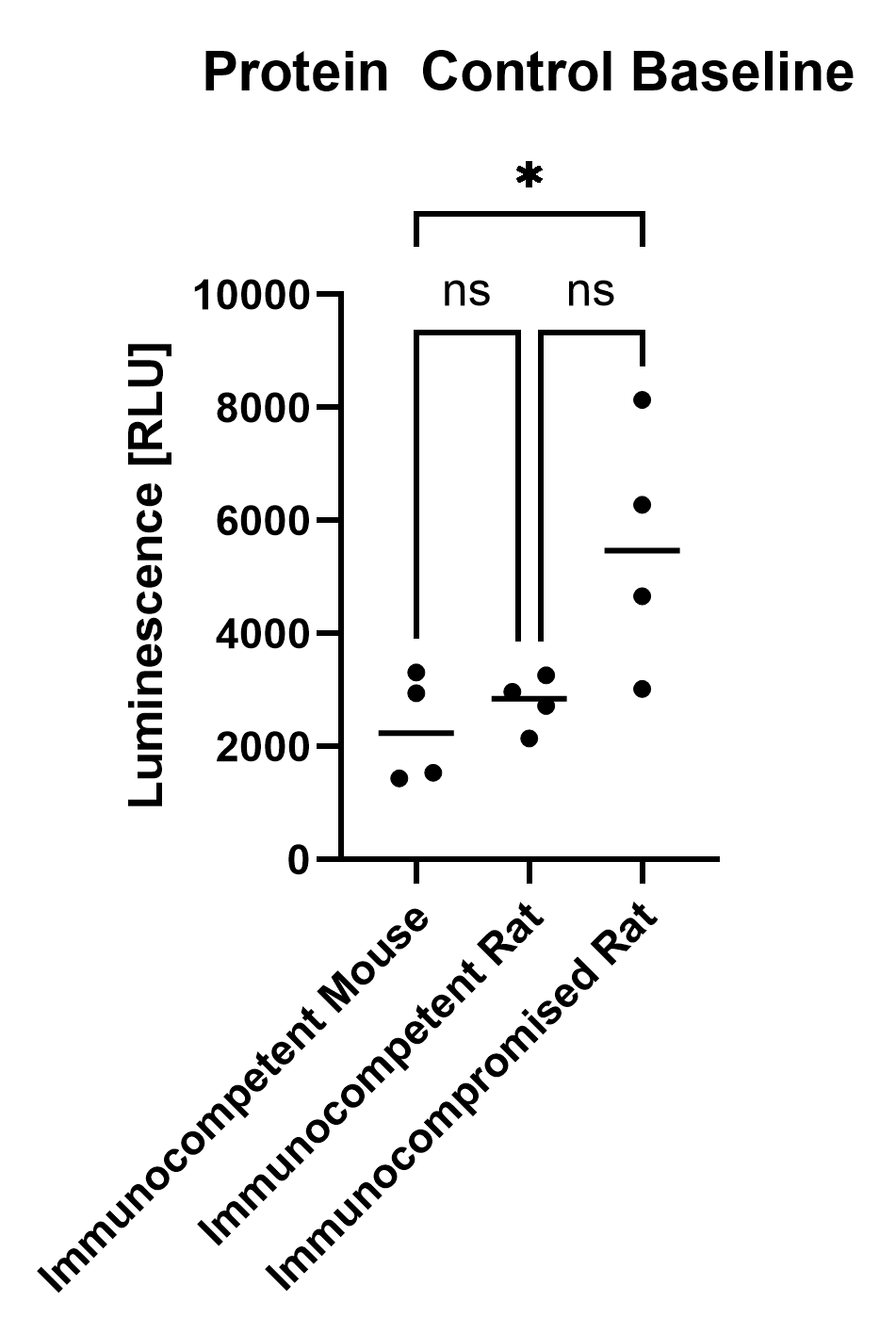

**Fig. S12.** Comparison of protein concentration baseline (pre-FUS sonication) for mouse, immunocompromised and immunocompetent rats.
